# From soil to sea: unravelling the metabolic versatility and social dynamics of Myxococcota bacteria from different Danish environments

**DOI:** 10.64898/2026.07.10.737566

**Authors:** Francesca Petriglieri, Yu Yang, Zivile Kondrotaite, Chenjing Jiang, Thomas Bygh Nymann Jensen, Mantas Sereika, Anders Ogechi Hostrup Daugberg, Kalinka Sand Knudsen, Francesco Delogu, Mads Albertsen, Caitlin Margaret Singleton, Per Halkjær Nielsen

## Abstract

Myxococcota are globally distributed bacteria renowned for their remarkable ecological and biotechnological significance due to their complex lifestyles, social behaviour, and secondary metabolite production. Despite their ubiquity in diverse environments, including soil, marine, and extreme habitats, their diversity and ecological roles remain underexplored. Here, we used the Microflora Danica dataset, encompassing >10,000 metagenomes and >400 rRNA gene datasets from various environments in Denmark, to investigate the distribution, diversity, and metabolic potential of Myxococcota. We show that Myxococcota are ubiquitous but their distribution is structured by environment, with soil-associated lineages enriched in predatory and multicellular development traits, whereas aquatic-associated taxa, particularly those from anoxic sediments, exhibit a higher potential of anaerobic metabolism and phototrophy. Comparative genomic analysis reveals widespread metabolic potential for secondary metabolite production, hydrocarbon degradation, and organohalide transformation, alongside predicted diverse contribution to carbon and nutrient cycling. Together, these findings propose Myxococcota as a functionally diverse and ecologically differentiated phylum, extending beyond canonical predation and multicellularity, and shows their promise as a reservoir of unexplored functional potential for biotechnological applications in drug discovery and environmental remediation.

## Introduction

Myxococcota are highly complex, socially coordinated bacteria with notable ecological and biotechnological importance [1]. Their hallmark traits include a predatory lifestyle and multicellular fruiting body formation triggered by nutrient limitation [1–3]. This complexity is exemplified by the model organism *Myxococcus xanthus,* which coordinates collective behaviours via adventurous (A) and social (S) gliding motility, the latter dependent on exopolysaccharide (ExoPS) production. These mechanisms enable cooperative predation through swarming, secretion of lytic enzymes and secondary metabolites, and efficient biomass degradation [1, 4]. Upon starvation, these same populations undergo a developmental program leading to fruiting body containing over 100,000 cells, differentiated in resistant myxospores, peripheral cells primed for rapid nutrient exploitation, and cells undergoing programmed cell death [3, 5, 6]. These transitions depend on fine-tuned differential gene expression, kin recognition, and sophisticated cell-to-cell communication, representing one of the most advanced forms of bacterial multicellularity [3, 5–8].

The biochemical capabilities underlying these behaviours also contribute to the remarkable metabolic diversity of Myxococcota. They use a wide range of antimicrobial molecules to inhibit and lyse competitors, many are targets in agriculture and medicine, particularly in antibiotic discovery [9–14]. These include ribosomally synthesized and post-translationally modified peptides (RiPPs), nonribosomal peptides (NRPs), polyketides (PKs), and terpenes, which can exhibit antimicrobial or cytotoxic activity [9, 13, 15, 16]. Their biosynthesis is typically encoded by biosynthetic gene clusters (BGCs) that coordinate precursor synthesis, enzymatic assembly, regulation, resistance, and transport [17–19]. As a result, Myxococcota are increasingly recognized as a valuable source of novel BGCs and bioactive compounds, especially in the context of rising antibiotic resistance [17, 20].

Importantly, their metabolic versatility also underpins a broad ecological distribution across diverse environments. Myxococcota have been isolated from soils, deserts, permafrost, contaminated sediments, and marine habitats [12, 21–29]. Culture-independent studies further show that they are widespread across soils, sediments, wastewater treatment systems, and even detected in the deepest oceans [5, 30–33]. Within these habitats, they contribute to biogeochemical cycling and organic matter turnover, with growing evidence of their involvement in processes relevant for bioremediation. Some species can reduce metals under anaerobic conditions and degrade aromatic hydrocarbons and xenobiotics [26]. Recent genome-resolved studies detected pathways for contaminant transformation and degradation of recalcitrant compounds, highlighting their potential for environmental remediation and sustainable applications [34].

Despite their ubiquity and biotechnological importance, the diversity and ecological functions of Myxococcota across environments remain poorly characterized. Here, we leverage the Microflora Danica (MFD) dataset [35, 36], comprising 10,683 short-read metagenomes, 154 long-read metagenomes, and >400 rRNA gene datasets, together with the Danish Microbial Database for Activated Sludge (MiDAS) genome project [32]. Using these datasets, we investigate the Myxococcota across diverse Danish habitats, with focus on the linkage between their phylogenetic diversity, habitat distribution, predatory traits, multicellular development, and broader roles in biogeochemical cycling and bioremediation.

## Materials and Methods

### Genome data collection and processing

A total of 215 medium-quality (MQ) and high-quality (HQ) dereplicated Danish Myxococcota species representatives (referring to both Myxococcota and Myxococcota_A phyla, hereafter, unless otherwise specified) were recovered as part of the MFD sequencing genome recovery effort [36] and the Danish MiDAS genome project [32] (SI1 Table S1). These genomes were retrieved from various kinds of soil, sediment, and wastewater treatment plants (WWTPs). Soil samples were mostly collected as topsoils at a depth of 0-20 cm except for the subterranean soils, where soils were collected around the oxic-anoxic interface. Detailed sampling strategies are found in the original publication [35]. Overall descriptions of sample metadata, habitat classification, and genome reconstruction approaches are provided in the supplementary material (SI2 Supplementary Methods). Genome completeness and contamination were assessed using CheckM1 [37] (v1.2.2) and CheckM2 [38] (v1.0.2). Quality was assigned based on the stringent minimum information about a metagenome-assembled genome (MIMAG) criteria [39] standards, including tRNA and rRNA gene presence. Taxonomy was classified based on GTDB-tk [40] (v2.4.1) using the conserved marker genes defined in GTDB [41] R226 release.

### Genome-based phylogenomic and 16S rRNA gene-based phylogenetic analyses

For genome-based phylogenomic analysis, all 215 HQ and MQ Danish Myxococcota genomes and 86 GTDB (release R226) [41] Myxococcota species representative genomes were used (SI1 Table S1). The 120 single-copy marker genes were identified from the Danish genomes and aligned with those in the GTDB-Tk [40] (v2.4.1) reference genomes using the *de novo* workflow. Fourteen *Bdellovibrio* genomes were used as an outgroup. The multiple sequence alignment was inputted to IQ-TREE [42] (v2.3.3) for tree generation with the WAG+G model and 1000x bootstrap iterations. The resulting tree was visualized in ARB v6.0.6 [43] to re-root using the selected outgroup and exported for visualization and final aesthetic adjustments in iTOL v6.1.1 [44] and Inkscape v0.92 (https://inkscape.org/).

The 16S rRNA gene sequences were extracted from the genomes using Fxtract v2.3 (https://github.com/ctSkennerton/fxtract). Subsequent phylogenetic analysis was performed using ARB v7.0, where a maximum likelihood phylogenetic tree was calculated with a 1000-replicates bootstrap analysis, based on comparative analysis of aligned 16S rRNA gene sequences retrieved from the Myxococcota genomes and the MFD database [35, 36].

### Statistical analyses

R [45] (v4.4.0) was used for statistical analysis. Significance of differences in the genome size for HQ Danish Myxococcota MAGs was statistically tested using a Kruskal–Wallis rank-sum test (function kruskal_test) and post hoc *Dunn* test for pairwise comparisons (function dunn_test) from rstatix [46] (v0.7.2).

### Screening of the distribution of Myxococcota in Denmark

A total of 10,752 Danish environmental samples (10,683 from MFD [35] and 69 from Danish MiDAS genome project [32]) were screened for the detection of Myxococcota across different habitats of soils, sediments, waters (including WWTPs). Microbial community composition was profiled using the “pipe” module of SingleM [47] (v0.20.3), with an expanded GTDB R226 metapackage v5.4.0 (DOI: https://zenodo.org/records/15232972) incorporating 17,716 HQ and MQ dereplicated Danish MAGs (both Illumina short-read [35] and Oxford Nanopore long-read [36] MFD metagenomes), added using the SingleM “supplement” module. Relative abundances were determined as lineage-specific coverage normalized to total coverages of all lineages per sample. After filtering low sequencing depth samples, 10,163 samples were retained for downstream analysis. Microbial profiles were visualized in heatmap using the R packages “ampvis2” v2.8.7 [48] and “ggplot2” v3.5.1 [49].

Genome abundances within all Danish metagenomes were estimated using sylph [50] (v0.6.1), with a reference database including species representatives from GTDB R226 and dereplicated MFD genomes from both short-read [35] and long-read [36] efforts. Samples and database genomes were sketched independently under the default parameters using the “sketch” subcommand. Genome abundances were then determined using the “profile” subcommand followed by the utility scripts released previously [51] to add sylph taxonomy (script sylph_to_taxprof.py) and extract quantification (script merge_sylph_taxprof.py).

Some observations of the environmental distribution of Myxococcota were also cross-compared to Sandpiper [47] (v2.1.1) which analyzes public metagenomes from NCBI SRA uploaded before 3^rd^ March 2026 using SingleM with taxonomy in GTDB R232. To limit to environmental habitats, samples from all host habitats were discarded. In addition, for samples within the “ecological” category, those with the following labels were removed: “Escherichia coli”; “viral metagenome”; “Sulfurovum”; “Spinacia oleracea”; “Equus caballus”; “Synchytrium endobioticum”; “Pseudomonadota”; “Acidobacteriota bacterium”; “Homo sapiens”; “Tetrao urogallus”; “synthetic metagenome”; and “bacterium”. The resulting total number of metagenomes analysized by Sandpiper [47] (v2.1.1) is 104,559.

### Genome mining analysis for identification of predatory machineries, fruiting body formation, and other metabolic pathways

For comparative genome mining, 90 HQ Danish Myxococcota MAGs and 11 HQ GTDB reference isolate genomes were annotated with KEGG orthology identifiers (KEGG FTP Release 2024-01-01) [52] using DRAM [53] (v1.4.6). KEGG module completeness was assessed with EnrichM (v0.6.5, github.com/geronimp/enrichM) ‘classify’, retaining modules with ≥80% completeness [54]. Genomes were additionally inspected in “MicroScope Microbial Genome Annotation & Analysis Platform” (MAGE) [55] to validate KO annotations and identify key genes (e.g., *SitA*) missed by automatic pipelines.

For assessing social behaviour potential, protein-coding genes were predicted and annotated using bakta [56] (v1.9.3). Predation- and fruiting body–related homologs were identified by Blastp [57] (pident >30%, e-value 1e-5, and qcovs >50% on bitscore-based best hits) against 208 *Myxococcus xanthus* reference proteins [5, 58] (SI1 Table S2). ExoPS production potential was expanded using epsSMASH [59] (v1.0, with default parameters). PSORTb 3.0 [60] (v3.0.2) was used to predict genes encoding extracellular enzymes. Biosynthesis pathways for bacteriochlorophyll and carotenoids were based on previously published metabolic schemes [31] (marker protein list in SI1 Table S3). Detection of *crtB* for carotenoids biosynthesis was expanded by including the Myxococcota *crtB* in uniprotKB with the accession A0A0K1E8V4 using blastp (≥30% ident, ≥75% qcovs, and ≤1e-20 e-value) [31].

Dissimilatory sulfate reduction was confirmed by verifying the phylogeny of DsrAB protein sequences as previously described [51]. Bioremediation potential related to hydrocarbon degradation was predicted using hmmer [61] (v3.4) for bakta-predicted genes against the CANT-HYD database [62] with the “noise” cutoff. Reductive dehalogenases were identified using blastp (pident ≥50% and qcovs ≥70% on bitscore-based best hits) on bakta-predicted genes against the Reductive Dehalogenase Database (RDaseDB) [63]. Polyethylene terephthalate (PET) hydrolase was predicted using PSORTb 3.0 [60] (v3.0.2) and manually verified by NCBI online blastp against the “refseq_protein” database (accessed on Sept 06 2025, top 5 hits were summarised in SI1 Table S4). Additionally, we assessed the phylogeny of these PET hydrolases with the 301 verified PET hydrolyzing enzymes downloaded from PAZy database [64]. The downloaded enzymes were dereplicated at 90% amino acid identity using usearch [65] (v12.0-beta) with the cluster_fast flag. Outgroups were chosen as hydroxyisourate hydrolases (AFQ59091, MDK2600047, MCY7886136) from the *Bacillus* genus as previously reported [66]. Verified sequences, our sequences, and outgroups werealigned using MAFFT [67] (v v7.525), and trimmed at a gap threshold of 0.2 using trimAl [68] (v1.5.rev1). Protein phylogeny was inferred using IQ-TREE [69] (v3.1.4) under the WAG model, with 1000 ultrafast bootstrap iterations.

### Genome mining analysis for identification of BGCs and potential antimicrobials

Secondary metabolite gene clusters were mined using AntiSMASH [70] (v7.1.0.1, with flags of “--cc-mibig --rre --fullhmmer --clusterhmmer --asf --cb-knownclusters --pfam2go”) with comparison to MiBIG [71] (v3.1) database. Detected BGCs were summarised using MultiSMASH (v.0.2.0) (Zenodo DOI: 10.5281/zenodo.8276143), and clustered into 8 major classes by BiG-SCAPE [72] (v1.1.9). To further assess biosynthetic potential for clinically important pharmaceutical drugs, these AntiSMASH-identified BGC regions were extracted using astool (https://github.com/BioGavin/astool) and submitted to the web-based classification tool NaPDoS2 [73] (with an additional cutoff of ≥50% aa identity to the default cutoffs). Antimicrobial-associated BGCs were identified based on the manually expanded annotation of the MiBIG [71] v3.1 database by searching activity-relevant keywords “antibacterial”, “antifungal”, “antiviral”, “antiprotozoal”, “antibiotic”, “antimalarial”, “antiparasidal”, and “anticoccidial” (SI1 Table S5).

### FISH probe design, optimization, and morphological characterization

FISH probes were designed using the ARB software v.6.0.6 [43]. Coverage and specificity were validated *in silico* with the Decipher web tool [74]. When needed, unlabelled competitors were designed. All probes were purchased from Biomers (Ulm, Germany), labelled with 6-carboxyfluorescein (6-Fam), indocarbocyanine (Cy3), or indodicarbocyanine (Cy5) fluorochromes. FISH was performed as described before [75]. The procedure for determination of the optimal formamide concentrations for each FISH probe and optimal hybridization conditions are detailed in SI2 Supplementary Methods and SI1 Table S6, respectively. EUBmix [76, 77] was used to target all bacteria. Microscopic analysis was performed with Axioskop epifluorescence microscope (Carl Zeiss, Germany) equipped with LEICA DFC7000 T CCD camera or a white light laser confocal microscope (Leica TCS SP8 X). Cell sizes were determined using ImageJ software [78].

### Naming of novel Myxococcota lineages

MAGs from novel genera with at most 10 contigs were named following the SeqCode rules [79] with the pipeline described previously [36]. Naming and explanation of the proposed names are provided in SI1 Table S7.

## Results and Discussion

### Phylogeny of novel Myxococcota from different Danish environments

A total of 215 Danish Myxococcota species representative genomes were recovered from the Microflora Danica (MFD) [35, 36] and Danish MiDAS [32] datasets, spanning 13 different ecosystem types (SI1 Table S8). Among the 90 HQ ones, genome sizes averaged 6.85Mbp (± 2.21Mbp, ranges 3.75Mbp – 11.5Mbp (SI1 Table S1). Interestingly, genome size distribution varied across habitats, with larger genomes predominantly associated with terrestrial and activated sludge environments (SI2 Figure S1). This pattern likely reflects adaptation to dynamic, nutrient-diverse conditions that favour metabolic versatility [80–82], whereas smaller genomes observed in sediment habitats may indicate reduced predatory and fruiting capabilities and more specialized or streamlined lifestyle [5, 80–82].

Phylogenomic analysis revealed that the 215 Danish MAGs were widely distributed across the Myxococcota phylum, encompassing a substantial fraction of uncultured diversity (Figure 1). In this study, eighteen HQ MAGs were assigned new genus and species names under SeqCode [79], and additional higher-rank taxa were proposed, including one class (from the phylum Myxococcota_A), two orders, and three families (SeqCode Registry Accession: seqco.de/r:yjqbk2e7) (SI1 Table S7). Based on GTDB-Tk classification and updated nomenclature, 134 MAGs were affiliated with the class Polyangiia, 49 with Myxococcia, three with Kuafuiibacteriia (former c WYAZ01), two each with the three additional novel classes with placeholder names UBA727, UBA796, and B64-G9, and one with Bradymonadia (genus *Persicimonas*). The remaining 22 MAGs belonged to the novel class Vahlivitia (named in this study in SI1 Table S7, former UBA9160, in Myxococcota_A). These phylogenomic assignments were consistent with patterns observed in 16S rRNA gene-based analyses performed using the MFD reference database (SI2 Figure S2), supporting the robustness of the recovered phylogenetic diversity.

**Figure 1.**
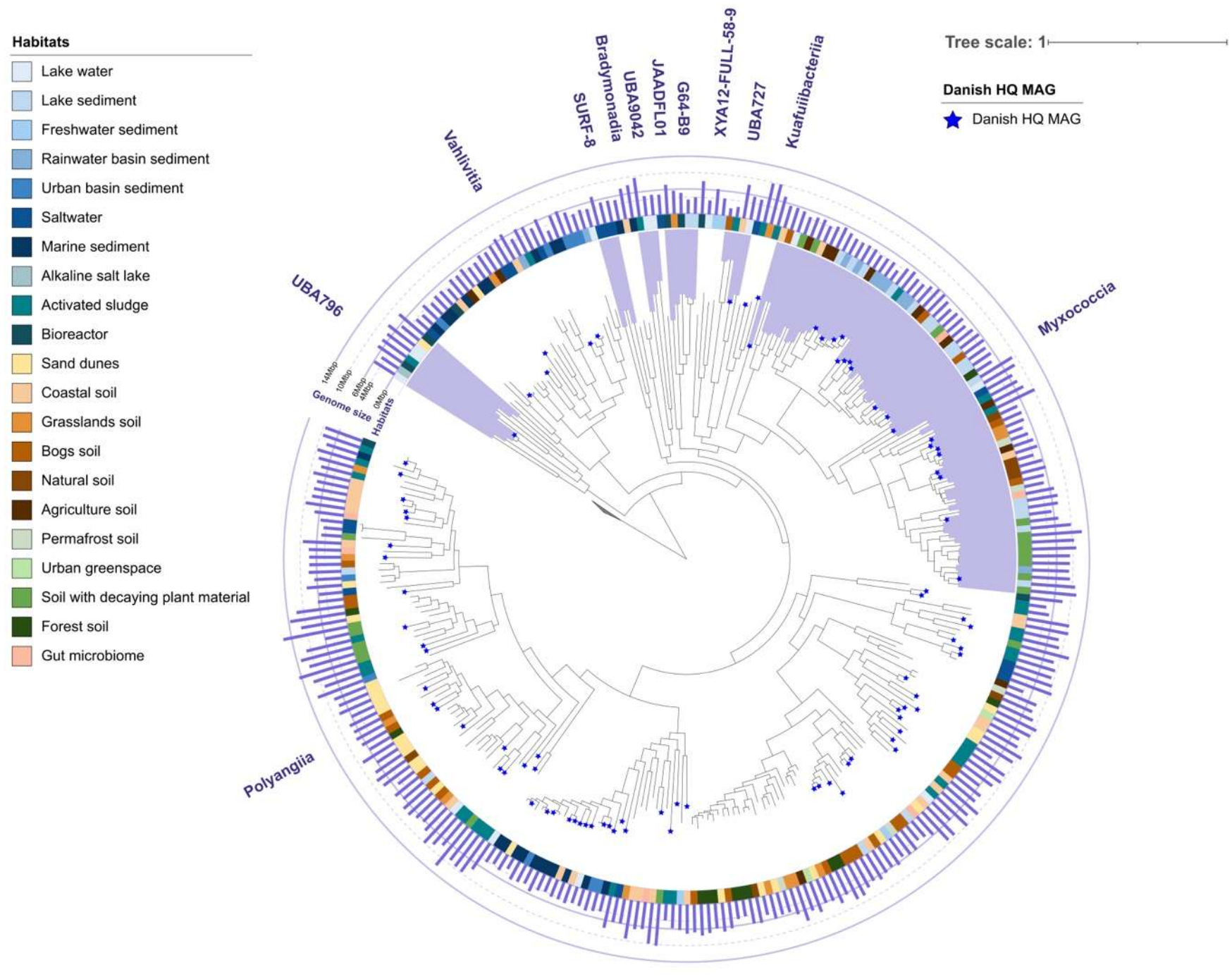
Phylogenetic genome tree showing the diversity of Myxococcota species representatives. The phylogenetic tree is based on the concatenated alignment of 120 single-copy marker gene proteins using GTDB-Tk. The 215 Danish species representative MAGs, retrieved from the MFD and the Danish MiDAS projects, were included in the tree together with 100 RefSeq representatives (GTDB R226 database [41]). Blue stars indicate HQ MAGs from different Danish environments. Additional information on the MAGs is presented in SI1 Table S1. Taxonomy at the class level is labelled in the tree with alternating white and purple shading for improved visibility. Annotations of the tree from the inner-most ring to the outermost ring are: 1) the different environments from which the MAGs were retrieved; 2) genome sizes, presented by the purple bar chart. The classes Vahlivitia and Kuafuiibacteriia were previously named as c UBA9160 and c WYAZ01, respectively. Classes UBA796 and Vahlivitia are from phylum Myxococcota_A and all other classes are from Myxococcota.

### Distribution of known and novel Myxococcota across different Danish environments

To determine the environmental distribution of Myxococcota, we profiled their occurrence across >10,000 Danish environmental samples [35] spanning diverse soil, sediment, and water habitats. Myxococcota were detected in all sampled habitats, demonstrating their ubiquity (Figure 2). However, their prevalence and community composition varied markedly across environments (Figure 2). Notably, they were highly abundant and consistently detected (≥1% relative abundance of microbial communities in >90% of samples) in natural saltwater sediments, agricultural soils, wetlands (bogs, mires and fens), and urban-associated soils (e.g., greenspaces, grasslands, roadsides, and landfill sites), identifying these as potential key ecological hotspots for Myxococcota (Figure 2A, SI1 Table S9). In contrast, some engineered systems (i.e., biogas plants, drinking water, and landfill leachates) and aqueous systems (i.e., groundwater and fjord waters) supported substantially lower abundances, indicating potential environmental constraints on their distribution. Across all habitats, Myxococcota typically constituted 0.2%-6.6% of the total microbial community, (Figure 2B), comparable to observations from the Earth Microbiome Project 16S rRNA gene datasets [30].

**Figure 2.**
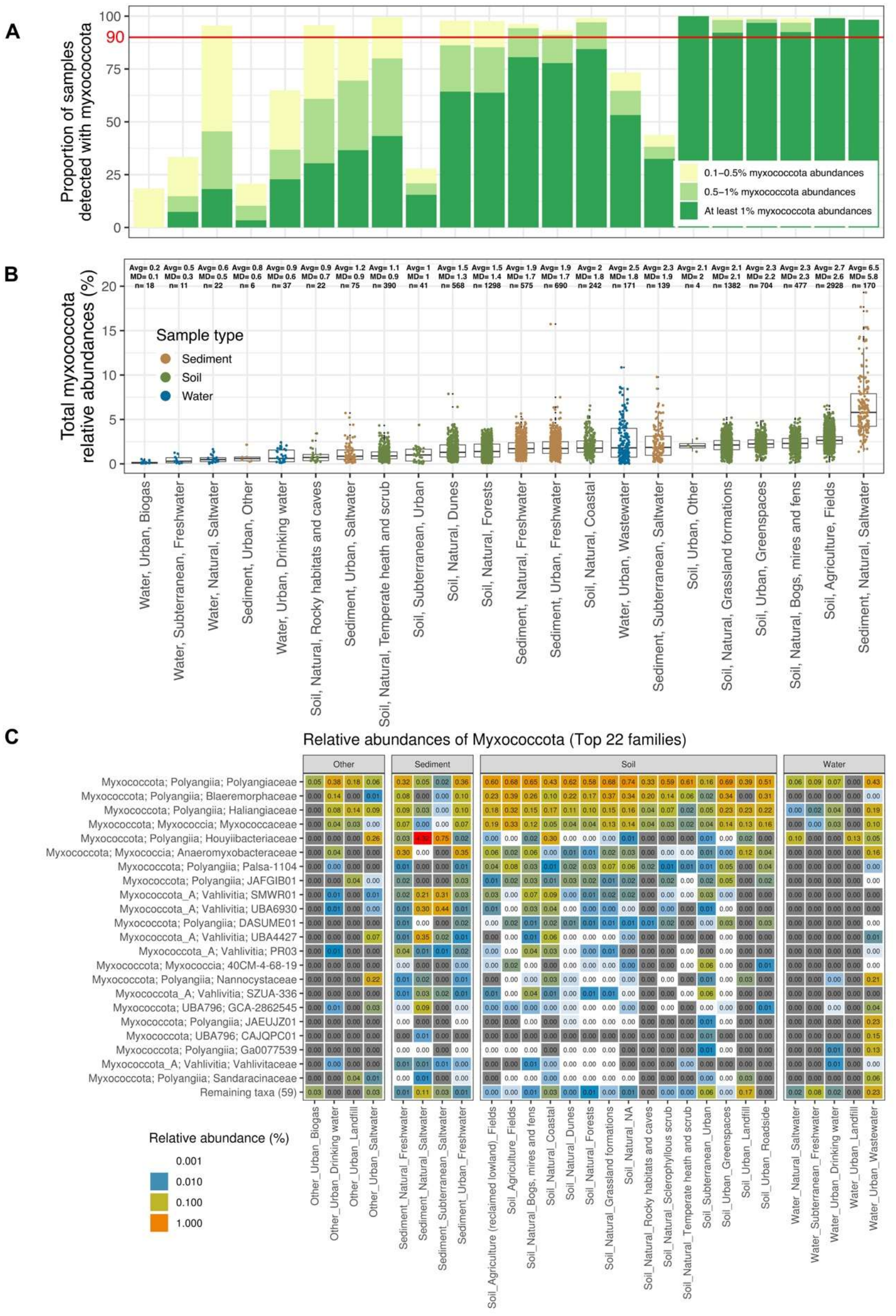
Detection, distribution, and abundance of Myxococcota across 10,163 Danish environmental samples spanning 28 habitats. **A)** Barplot showing the proportion of samples with Myxococcota detected in each habitat in 3 different Myxoccoccota prevalence categories. Note that Water_Subterranean_Freshwater and Water_Natural_Saltwater refer to groundwater and fjord waters, respectively; **B)** Boxplots showing the distribution of total Myxococcota abundances per sample (dots) across 28 habitats; **C** Heatmap showing the average relative abundances of the top 22 Myxococcota families (taxonomy in phylum;class;family) across 28 Danish environmental habitats (habitat label showing the sample type, area type, and MFDO1). Values in the heatmap represent the average relative abundances within the respective habitat. Values in the brackets along the x axis refer to the number of samples within each habitat that have Myxococcota detected. Avg: average, MD: median, N: sample number. Vahlivitia: former c UBA9160; Blaeremorphales: former o Fen-1088; *Vahlivitaceae*: former f UBA9160; *Houyiibacteriaceae*: former SG8-38; *Blaeremorphaceae*: former f Fen-1088.

Habitat preferences were further resolved at finer taxonomic resolution using single copy marker genes-(SingleM, Figure 2C) and k-mer-based (sylph, SI2 Figure S4) approaches, revealing pronounced ecological specialization among lineages. For example, the families *Houyiibacteriaceae* (former SG8-38, from class Polyangiia) and UBA6930 (from the class Vahlivitia), were strongly associated with saline environments, including saltwater sediments and waters, and coastal soils and rare in other soils and non-saline habitats, whereas *Myxococcaceae*, *Haliangiaceae*, *Anaeromyxobacteraceae*, and *Blaeremorphaceae* (former Fen-1088) were scarce in saline habitats (Figure 2C, SI2 Figure S4). Most notably, the uncultivated genus *Thyboronia* (former g SG8-38 in the family of *Houyiibacteriaceae*) were highly abundant (2.56% average, up to 14.07% based on sylph) in all MFD saltwater sediment samples. When broadening our habitat search for these families using Sandpiper [47], similar patterns were also observed (SI2 Figure S5, SI1 Table S10). Within non-saline environments, distinct niche partitioning was also evident: *Anaeromyxobacteraceae* preferentially occupied freshwater environments, while *Myxococcaceae*, *Blaeremorphaceae*, and *Polyangiaceae* exhibited a broad ecological distribution across various soils, with some lineages from these families showing high abundances in wetland habitats (Figure 2C, SI2 Figure S4). WWTPs hosted a distinct assemblage, including both well-characterized (e.g., *Polyangiaceae*, *Nannocystaceae*, *Haliangiaceae*, *Anaeromyxobacteraceae*, and *Myxococcaceae)* and uncharacterized families across different classes, highlighting engineered systems as reservoirs of unique Myxococcota diversity. Collectively, these results demonstrate that while Myxococcota are globally distributed, their distribution is strongly structured by environmental factors, such as salinity [83], resulting in clear habitat specialization across lineages. Interestingly, a substantial fraction of detected diversity lacked representative HQ genomes and could not be resolved taxonomically (SI2 Figure S6, SI1 Table S11), emphasizing the extent of the remaining unexplored diversity within this phylum.

### Potential for predation machineries in Myxococcota from diverse environments

We next investigated whether the observed ecological differences are reflected in the genomic potential for predation and social behaviour, a defining feature of the phylum [1]. While predation is traditionally considered a universal trait, emerging evidence suggests it may primarily represent an adaptation to terrestrial ecosystems, with non-soil lineages exhibiting alternative lifestyles [5]. To investigate this, we assessed key pathways underlying motility, chemosensory systems, secretion, production of ExoPS, hydrolytic activity, and secondary metabolites across 90 Danish HQ Myxococcota MAGs and 11 reference genomes from GTDB (see SI2 Supplementary Note for details).

Motility occurs in *M. xanthus* via A-motility and S-motility. A-motility involves proton motive force-driven gliding via the AglRQS motor complex (MotA/MotB homologs), linked to MreB cytoplasmic filaments [84–86]. Homologs of key A-motility genes were broadly distributed in the class Myxococcia (only exception of some *Anaeromyxobacteraceae* MAGs), Kuafuiibacteriia, and UBA727 (Figure 3). A few MAGs in *Blaeremorphaceae* and *Houyiibacteriaceae* also encoded homologs of key genes for A-motility (Figure 3). This feature did not appear to be ecosystem-specific, occurring in both soil- and freshwater-associated taxa (Figure 3, SI2 Figure S4). When these gene sets were partial or absent, we hypothesize that these lineages rely solely on S-motility, similar to observations from some isolates [26]. In contrast, S-motility relies on type IV pili (Pil complex), ExoPS secretion, and a specialized chemosensory network [85, 87]. Type IV pilus machinery homologs were nearly ubiquitous, with few exceptions in *Houyiibacteriaceae*, *Sandaracinaceae*, and *Nannocystaceae* (Figure 3). However, the prediction of type IV pilus assembly is not an unambiguous indication for S-motility potential, as these cell structures may be involved in other processes, such as adhesion, secretion, and uptake, or virulence [84, 88]. Consistent with this, ExoPS biosynthesis genes (*eps* gene clusters) and key chemosensory network components (Dif and Frz) were largely restricted to the class Myxococcia, whereas the regulators MglAB and RomR were broadly conserved and present in at least 90% of HQ Danish MAGs (Figure 3, Supplementary Data 1). Notably, Frz homologs were absent from the family *Haliangiaceae*, including *Haliangium ochraceum*, despite experimental evidence for S-motility in the isolate [24]. This suggests the potential use of alternative coordination mechanisms, likely involving only homologs of the chemotaxis system Dif, which is more widespread throughout the phylum (Figure 3).

**Figure 3.**
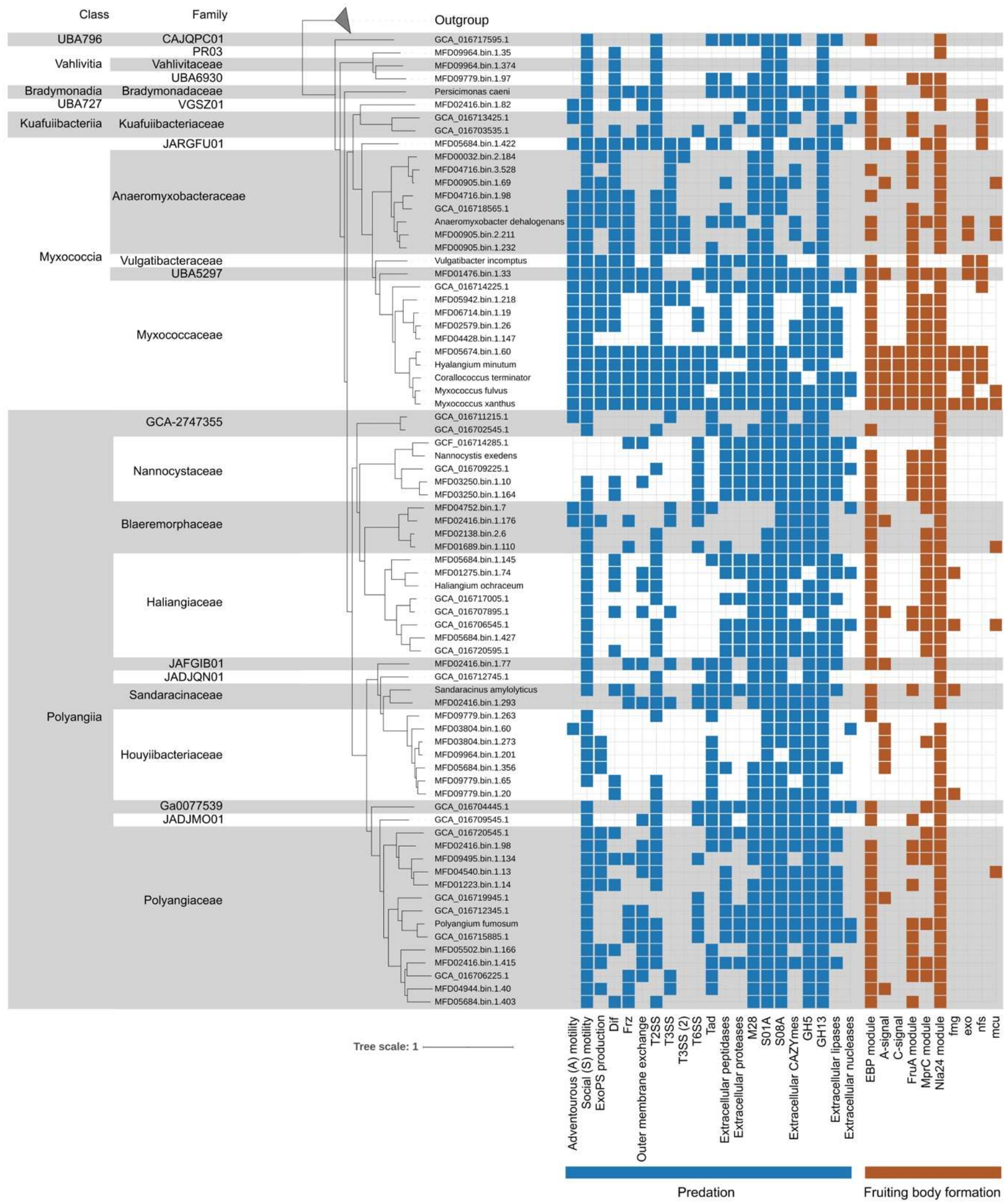
Predatory and fruiting body formation machineries of a subset of Myxococcota MAGs. Phylogenomic tree of subsetted Myxococcota genomes from GTDB and Denmark. Genomes are separated by shades and taxonomically labelled at the family level on the left. Vahlivitia: prev. c UBA9160; Kuafuiibacteriia; prev. c WYAZ01; Blaeremorphales: prev. o Fen-1088; *Kuafuiibacteriaceae*; prev. f WYAZ01; *Vahlivitaceae*: prev. f UBA9160; *Houyiibacteriaceae*: prev. SG8-38; *Blaeremorphaceae*: prev. f Fen-1088. Classes UBA796 and Vahlivitia are from phylum Myxococcota_A, and all the rest classes are from phylum Myxococcota.

Beyond motility, social interaction is further mediated by outer membrane exchange (OME), enabling resource sharing and kin recognition via TraA-TraB mediated transfer of membrane components and diverse SitA toxins, with immunity conferred by the co-expressed *sitI* gene [5, 8, 89]. In our dataset, TraB homologs were widespread (85/90 HQ MAGs), but TraA (29/90 HQ MAGs) was restricted to selected lineages in Myxococcia, UBA727, Kuafuiibacteriia, Bradymonadia, and Polyangiia (Figure 3, Supplementary Data 1). Toxin-associated SitA genes were restricted to *Myxococcus* relatives, in line with the fact that toxin-based OME is lineage-specific [5, 8, 89] (Figure 3). Together, these patterns indicate that canonical social and predatory coordination systems are unevenly distributed and largely enriched in soil-associated clades, reflecting evolutionary diversification in non-terrestrial lineages.

Predatory capacity also depends on extracellular killing and nutrient acquisition. *M. xanthus* lyses prey using secreted hydrolytic enzymes, such as amidases, glucosaminidases, and β-1,6-glucanases, and secondary metabolites targeting cells walls’ macromolecules [1, 27, 90, 91]. Subcellular localization analysis of predicted proteins in the MAGs revealed widespread occurrence of extracellular peptidases/proteases, with few exceptions in *Polyangiaceae*, *Houyiibacteriaceae*, *Blaeremorphaceae*, *Anaeromyxobacteraceae*, and in the novel class UBA727 (Figure 3). Most predicted extracellular peptidases belonged to MEROPS families M28, S8, and S1 (Figure 3, Supplementary Data 2), encompassing enzymes capable of peptidoglycan cleavage, bacteriocin activity, and extracellular protein digestion [5]. Extracellular carbohydrate-active enzymes (CAZymes) were also prevalent, particularly glycoside hydrolases GH5 (cellulose degradation), GH13 (peptidoglycan lyases and amylases), and polysaccharide lyases (pectin degradation) (Figure 3, Supplementary Data 3), consistent with the ability to degrade both extracellular polysaccharides and cellular substrates, thus functioning as direct prey-killing factors. These enzymes were in some cases complemented by lipases and nucleases, likely contributing to full prey biomass use (Figure 3, Supplementary Data 3). These degradative capabilities were supported by multiple secretion systems: the type II secretion system (T2SS) was widespread (except for some *Nannocystaceae* and *Anaeromyxobacteraceae*) (Figure 3, Supplementary Data 1), whereas type III secretion systems (T3SS) and T3SS(2), known in *M. xanthus* to mediate enzyme export [88, 92, 93], were primarily restricted to the class Myxococcia (Figure 3, Supplementary Data 1). Type VI secretion systems (T6SS) and Tad-like machineries were more broadly distributed across the phylum (Figure 3, Supplementary Data 1). While experimental evidence has shown that mutations of the T6SS does not impair *M. xanthus* predation abilities and it is unlikely involved in predation [88], T6SS may support contact-dependent predatory mechanisms in other Myxococcota lineages, suggesting functional diversification in enzyme delivery and interaction strategies across lineages.

Consistent with their competitive lifestyle, Myxococcota also exhibit extensive biosynthetic potential, including pigments, siderophores, bacteriocins, and antimicrobials [9]. AntiSMASH detected a total of 918 BGCs from these 90 HQ Danish Myxococcota MAGs (SI1 Table S12), of which 89.4% were complete. The highest BGC counts per HQ MAG were observed in the families *Haliangiaceae* (27 BGCs) and *Myxococcaceae* (23 BGCs) (Figure 1, SI1 Table S12, SI2 Figure S7), identifying these as major reservoirs of secondary metabolite diversity. Domain-based analysis using NaPDoS2 [73] detected 55 ketosynthase and condensation domains linked to 26 known antimicrobial compound classes, such as stigmatellin, chondrochloren, and myxalamid, inhibiting a broad range of bacteria [94] (Figure 4). Overall, homology (53-67% aa identity, 10^th^-90^th^ percentile) to characterized domains suggests both conservation and diversification of antimicrobial biosynthesis, supporting a role for chemical-mediated predation and competition.

**Figure 4.**
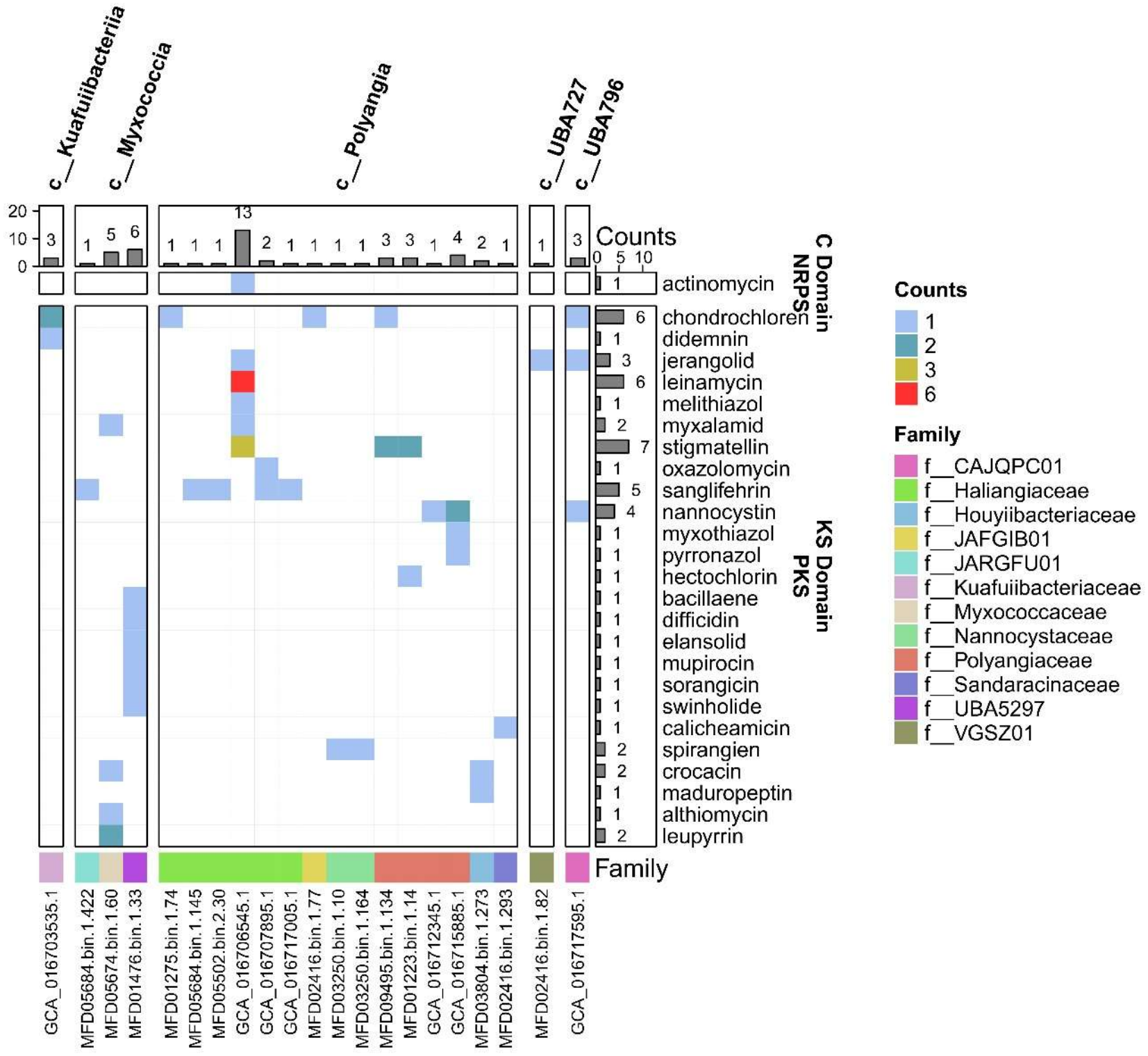
Biosynthetic potential for antimicrobial secondary metabolites in Danish Myxococcota species representatives. Heatmap showing the counts of identified genes for the production of antimicrobial secondary metabolites (C domain for NRPS and KS domain for KPS) in Danish Myxococcota. The barplot on top summarises the total domain sequence counts per genome, and the barplot on the right summarises the total domain sequence counts per compound. *Houyiibacteriaceae*: prev. SG8-38; Class UBA796 is from phylum Myxococcota_A, all other classes are from phylum Myxococcota.

Integrating all these traits reveals a potential ecological signal: soil-associated lineages possess the well-characterized, canonical *M. xanthus*-like complete predatory and social toolkits (notably *Myxococcaceae* and *Polyangiaceae*), whereas aquatic lineages from marine, freshwater, and sediment environments (*Houyiibacteriaceae*, *Anaeromyxobacteraceae*, Vahlivitia) frequently lack these typical markers, suggesting they have evolved fundamentally divergent or currently uncharacterized coordination mechanisms. This pattern supports the hypothesis that complex predation and social behaviour evolved primarily as an adaptation to nutrient-variable terrestrial ecosystems, while non-soil Myxococcota rely on alternative ecological strategies, underscoring a broader functional and evolutionary diversification within the phylum than previously recognized [5].

### Cellular differentiation and sporulation mechanisms in Myxococcota from different environments

Given the uneven distribution of known predatory traits across habitats, we next examined the genomic potential for multicellular development, a defining but unevenly distributed feature of Myxococcota. In *M. xanthus*, starvation triggers coordinated aggregation and fruiting body formation via complex regulatory cascades involving transcriptional modules (e.g., EBP, FruA, Nla regulators) and ExoPS-mediated cell cohesion [6, 7, 95, 96]. Homologs of these regulators were widespread in the MAGs (85.6%, 63.3%, 100.0%, 83.3%, 100.0% of the 90 HQ Danish Myxococcota for Nla18, Nla28, Nla6, Nla24, and dmxB, respectively) (Figure 3, Supplementary Data 1), although their broad occurrence in bacterial genomes limits their predictive value [5]. Co-occurrence of ExoPS genes and early-stage aggregation transcription factors provide a more robust indicator of developmental potential and was predominantly observed in soil-associated lineages (*Myxococcaceae*, Figure 3, SI2 Figure S4). Finally, the Mrp module (MrpABC), which cooperates with FruA to regulate aggregation and sporulation [5–7, 21], was detected primarily in *Myxococcaceae*, *Haliangiaceae*, and *Nannocystaceae* families. Many lineages lacked complete regulatory modules, suggesting limited or absent capacity for canonical fruiting body formation (Figure 3, Supplementary Data 1) [5]. Similarly, genes associated with sporulation in *M. xanthus* (*fmg*, *exo*, *mcu*, *nfs,* and spore coat protein genes), were fully conserved in *Myxococcaceae,* with partial or sporadic representation in other families (*Polyangiaceae*, *Nannocystaceae*, *Haliangiaceae,* and *Kuafuiibacteriaceae*). However, the prediction of sporulation based solely on gene presence is hindered by their involvement in other core cellular processes, including cell cycle regulation and division [97]. Notably, some known spore-formers (e.g., *H. ochraceum*, *N. exedens*) [28, 29, 97], lack canonical gene sets (Figure 3), indicating that alternative, lineage-specific mechanisms for multicellular development likely exist beyond well-characterized models.

To complement genomic predictions, we designed two FISH probes (SI1 Table S6, SI2 Figure S2) targeting the class Polyangiia and the genus UBA4427 (class Vahlivitia) in environmental samples. In activated sludge, the class Polyangiia exhibited diverse morphologies, including rod-shaped cells in large circular clusters (Fig 5A, Fig 5F, SI1 Table S15), fibre-like structures (Figure 5C, SI1 Table S15), and single or clustered rods (Fig 5D-E, SI1 Table S15), while UBA4427 bacteria were rod-shaped and mostly found in small clusters (Figure 5B). Some of these spatial arrangements are consistent with facultative social behaviour and potential predatory behaviour interaction *in situ.* The arrangements likely influence bacterial population sizes, metabolize cellular debris, and support nutrient removal, with some genera (e.g., *Haliangium*) potentially contributing to both phosphorus and nitrogen cycling [64, 98]. However, while the observed morphologies are consistent with potential sociality for the class Polyangiia, they could also likely be explained by the nature of the matrix in the WWTP environment (i.e., the physical and chemical architecture). Therefore, time-lapse or targeted cultivation experiments are required to definitively link these spatial arrangements to active predation in the future.

**Figure 5:**
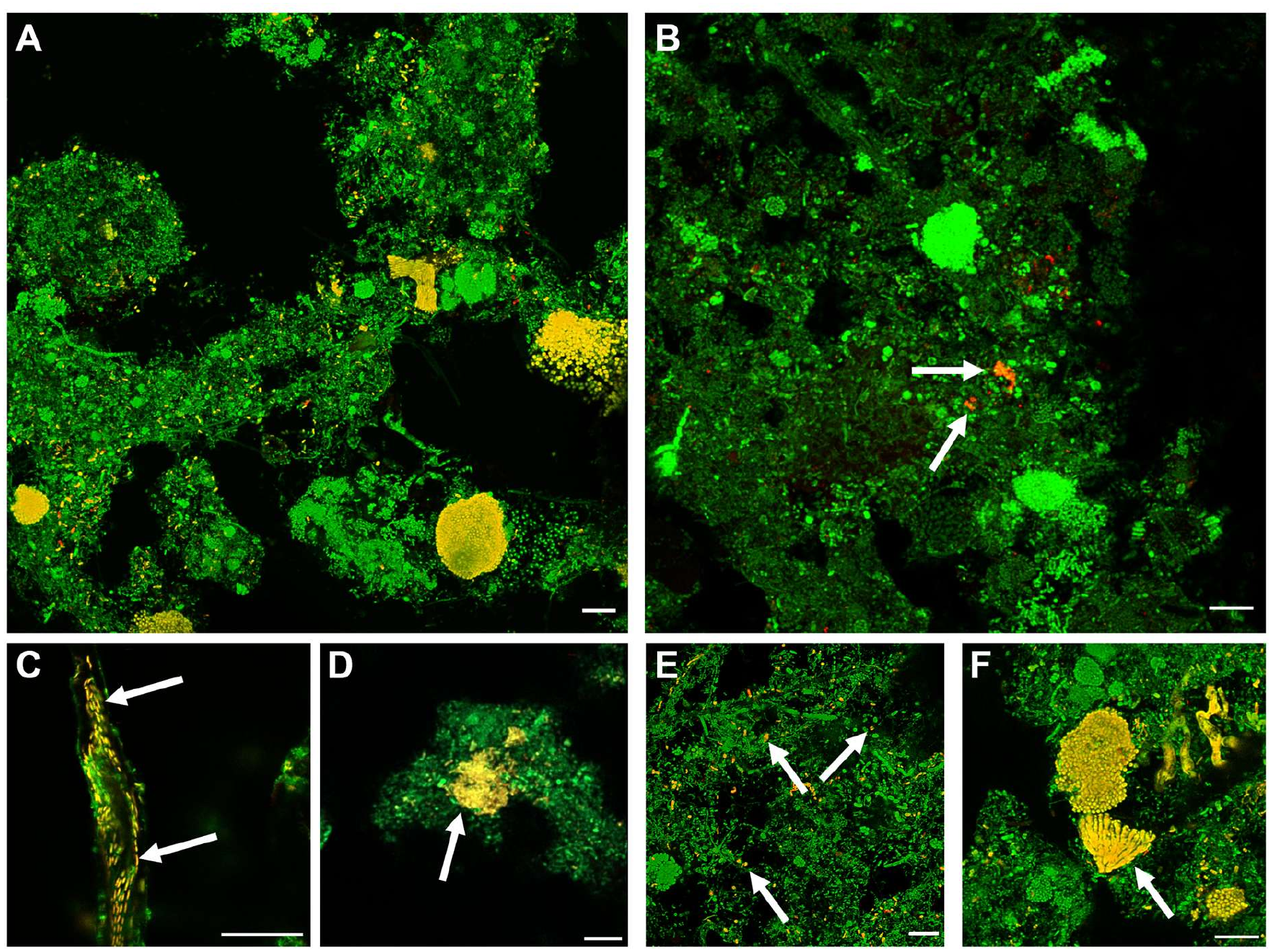
FISH micrographs of different morphologies of Myxococcota cells in full-scale activated sludge from WWTPs. All bacteria were targeted with EUBmix (green). Specific probes in **A)** and **B)** target Polyangiia class (Myxo493, yellow) and UBA4427 class in Myxococcota_A phylum (Myxo808, orange), respectively. FISH **C)-F)** images represent different morphologies and spatial arrangement within the Polyangiia class. The scale bar is 10 µm in all images. White arrows in **B)-F)** images indicate cells positive for the specific probe.

### Myxococcota possess a versatile lifestyle in different environments

Functional profiling revealed a highly versatile and habitat-linked metabolic repertoire across Myxococcota, consistent with their broad environmental distribution. They encoded potential for a heterotrophic lifestyle, with complete or near-complete central carbon metabolism, (glycolysis, pentose phosphate pathway, TCA cycle), alongside extensive carbohydrate degradation potential supported by diverse glycoside hydrolases and polysaccharide-degrading enzymes and sugar transporters (*msmX*, *thu*, *frc*, *xyl*) (Figure 6, Supplementary Data 3). Similarly to cultured representatives [5, 22, 100, 101], peptides and amino acids appeared to be key substrates in the Danish MAGs, as indicated by widespread transporters (*opp*, *liv*, *glt*), and catabolic pathways (Figure 6, Supplementary Data 4,5). In addition, several MAGs encoded potential for beta-oxidation of long-chain fatty acid (56/90), acylglycerol degradation (18/90), acetate utilization through the phosphate acetyltransferase-acetate kinase pathway (*actP* for uptake, *acs*, *pta,* and *ackA* for use, 38/90), and a complete glyoxylate cycle (39/90) (Figure 6, Supplementary Data 5), highlighting metabolic flexibility that likely supports survival under fluctuating nutrient availability, for example in soils or activated sludge systems.

**Figure 6.**
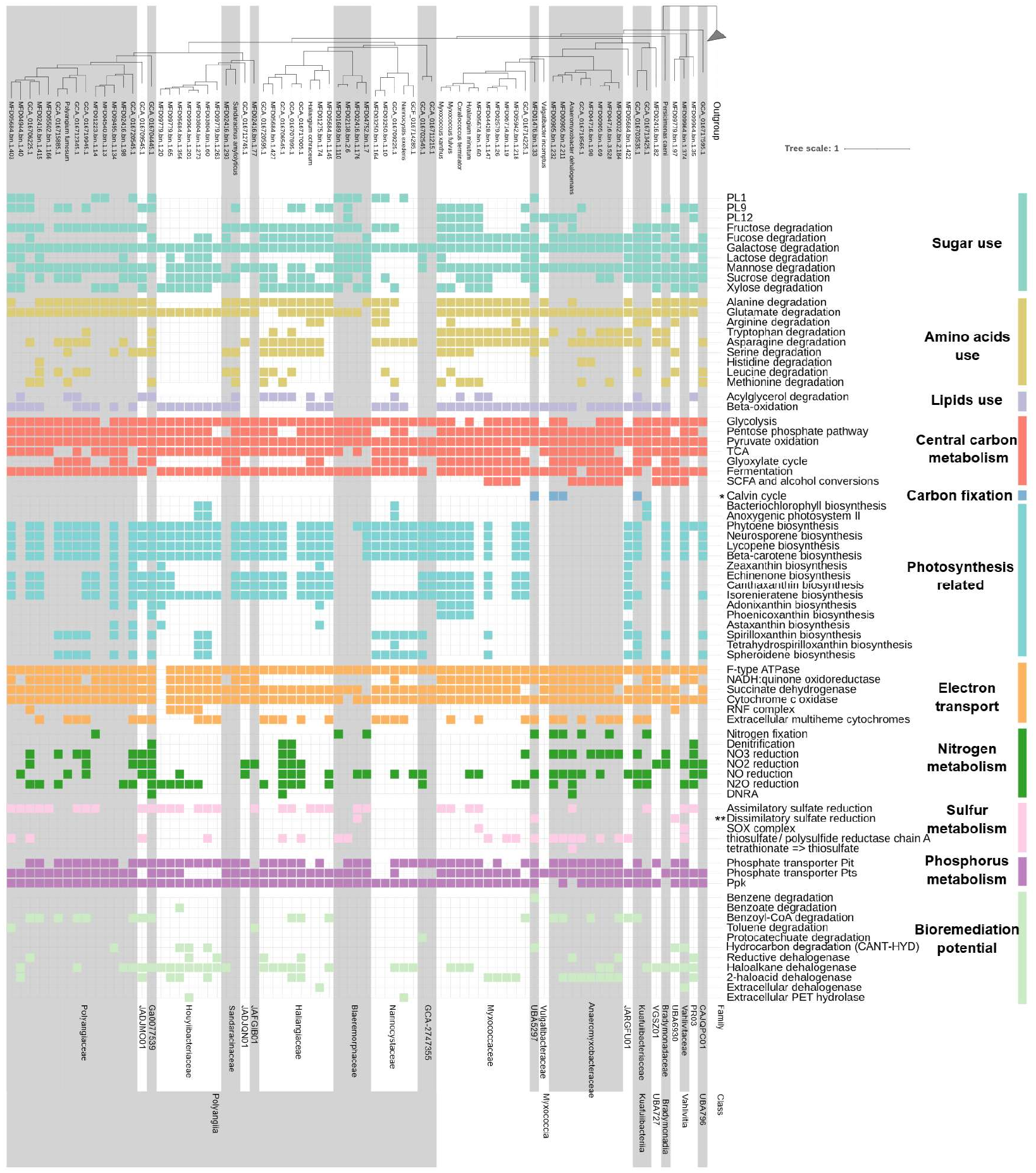
Metabolic and bioremediation potential the Myxococcota MAG subset. Phylogenomic tree of subsetted myxococcota genomes from GTDB and Denmark. Genomes are separated by shades and taxonomically labelled at the family level on the left. Presence/absence of the functional potential is indicated by at least 80% module completeness in KEGG. Asterisks denote if the presence of key enzymes were checked (details in Supplementary Data 2): * means present with KEGG K01601 and an optional presence of K01602. These two represent RuBisCO large and small subunits, respectively.), and ** means present with both KEGG K11180 and K11181 for dsrA and dsrB, respectively). Vahlivitia: prev. c UBA9160; Kuafuiibacteriia; prev. c WYAZ01; *Kuafuiibacteriaceae*; prev. f WYAZ01; *Vahlivitaceae*: prev. f UBA9160; *Houyiibacteriaceae*: prev. SG8-38; *Blaeremorphaceae*: prev. f Fen-1088. Classes UBA796 and Vahlivitia are from phylum Myxococcota_A, and all the rest classes are from phylum Myxococcota.

Beyond heterotrophy, a subset of lineages exhibited alternative metabolic strategies that may be linked to specific habitats. Phototrophic potential, including full bacteriochlorophyll biosynthesis and light harvesting pathways, was identified in 4 MAGs (in *Kuafuiibacteriaceae*, *Nannocystaceae*, and *Houyiibacteriaceae*) retrieved from activated sludge, coastal sediments or bogs soils, further expanding the range of families with potential phototrophic capability (Figure 6, SI2 Figure S4, SI1 Table S16). Several MAGs, spanning *Nannocystaceae*, *Anaeromyxobacteraceae*, *Houyiibacteriaceae,* and *Kuafuiibacteriaceae*, encoded a complete or near-complete Calvin–Benson–Bassham cycle including type I RuBisCO (4/90). However, only *Kuafuiibacteriaceae* encodes RuBisCO in additional to phototrophy genes, indicating it is a potential photoautotroph [31], while the other 3 putative phototrophic populations are potentially photoheterotrophs [31] (Figure 6). Carotenoids biosynthesis was widespread across Myxococcota, both in terrestrial and aquatic habitats (SI2 Figure S4), with over half of the MAGs (56/90) encoding at least one carotenoid biosynthesis pathway, most commonly phytoene (58/90), beta-carotene (52/90), neurosporene (52/90), and lycopene (52/90) (Figure 6, SI1 Table S17), likely supporting phototrophy, oxidative stress protection, and possibly antimicrobial activity [5, 31]. Together, these traits suggest that light-driven energy acquisition and photoprotection may provide adaptive advantages in stratified or nutrient-limited environments, including sediments and engineered systems.

While most cultured Myxococcota are aerobic, several species also exhibited the potential for facultative anaerobic metabolism, consistent with their presence in redox-variable environments (Figure 6, Supplementary Data 3,5). Fermentative pathways (to ethanol or lactate), formate dehydrogenase (*fdh*), and alternative respiratory systems were detected across multiple lineages (Figure 6, Supplementary Data 3,5). While nearly all genomes encoded cytochrome c oxidase genes (*cox* and/or *coo*), the few exceptions (in *Myxococcaceae*, *Nannocystaceae*, *Houyiibacteriaceae*, *Polyangiaceae*, *Anaeromyxobacteraceae*, *Vahlivitaceae,* and *Blaeremorphaceae* that were mostly from sediment- and water-associated habitats) likely relied on high affinity cytochrome *bd* ubiquinol oxidases (Supplementary Data 3), potentially detoxifying traces of O_2_ present under anaerobic conditions [5]. Some MAGs contained genes for additional energy-conserving systems, such as the *Rhodobacter* nitrogen fixation (RNF) complex and HydABC electron-bifurcating hydrogenases (Figure 6, Supplementary Data 3), which may contribute to proton motive force generation and ATP synthesis in anaerobic environments and further indicate adaptation to fluctuating oxygen availability in aquatic and sediment environments [5].

Further investigation into alternative electron acceptors indicated roles in biogeochemical cycling across habitats. Genes for nitrate reduction (*narGHI* or *napAB*) and nitrite reduction (*nirK/nirS*) were widespread, with some MAGs additionally encoding nitric oxide reduction (*norBC*) and nitrous oxide reduction (*nosZ*) (Figure 6, Supplementary Data 3). Complete denitrification potential was rare (4/90 MAGs) and largely confined to specific environments such as saltwater habitats (Vahlivitia), freshwater sediments (*Anaeromyxobacter*), natural soils (*Polyangiaceae*), and activated sludge (f Ga0077539, *Polyangiaceae*, *Haliangiaceae*) (SI2 Figure S4). Dissimilatory nitrate reduction to ammonia (DNRA) via the cytochrome-linked nitrite reductase NrfAH and nitrogen fixation were infrequent (Figure 6, Supplementary Data 3). Within the phylum, nitrogen cycling is well-characterized in *Anaeromyxobacter dehalogenans* [31, 102, 104] and suggested for abundant *Haliangium* species in activated sludge [105] but remains poorly described in most lineages. Phosphorus acquisition (Pit, Pst) and polyphosphate (polyP) metabolism were broadly conserved (Figure 6, Supplementary Data 3). In *M. xanthus*, polyP plays a significant role in supplying energy for motility, predation, and maturation of spores [105]. These findings indicate that Myxococcota may contribute to nitrogen and phosphorus turnover in diverse habitats, with particular relevance for aquatic ecosystems under anthropogenic influence, such as eutrophication [107].

Potential involvement in sulfur cycling was also detected, including dissimilatory sulfate reduction (6/90 MAGs, from *Blaeremorphaceae*, UBA5297, and UBA6930) and thiosulfate reduction (27/90) (Figure 6, Supplementary Data 3). The reductive nature of DsrAB was confirmed through phylogenetic analysis [51] (SI2 Figure S8). A few MAGs (from *Vahlivitaceae* and *Haliangium*) also encoded the Sox (sulfur oxidation) enzyme complex (Figure 6, Supplementary Data 3), which could facilitate both mixotrophic growth in oxygen-limited niches and contribute to the regeneration of sulfate for other microorganisms, reinforcing their ecological role in coupled carbon–sulfur cycles in anoxic niches such as sediments and wetlands [108]. Differences in sampling depth should be considered when comparing anaerobic metabolic potential between soil and aquatic sediment samples, as it may introduce biases. Our soil samples represented the upper 0–20 cm soil layer, and therefore may not fully capture the deeper anaerobic soil microhabitats. However, the presence of complete denitrification potential in natural soil-derived *Polyangiaceae* suggests that anaerobic or low-oxygen metabolic strategies can also occur within our sampled soil depth.

Notably, several lineages exhibited functions associated with environmental remediation processes. Reductive dehalogenases are well-described in *A. dehalogenans* [31, 109] and several homologs were identified in 8 HQ Danish MAGs across multiple families (*Anaeromyxobacteraceae*, *Haliangiaceae, Houyiibacteriaceae, Polyangiaceae,* and class Vahlivitia), particularly in lineages associated with sediments, soils, and wastewater systems (Figure 6, SI1 Table S18, Supplementary Data 6). Alongside respiratory dehalogenation, the MAGs also encoded non-respiratory dehalogenases (e.g., haloalkane dehalogenases or 2-haloacid dehalogenases) (Figure 6, SI2 Table S19), enabling both energy-conserving and non-energy-conserving halogen removal, suggesting that Myxococcota are versatile mediators of organohalide transformation, making them promising targets for bioremediation of contaminated soils, sediments, and wastewater systems [110, 111]. Similarly, genes involved in hydrocarbon degradation pathways (e.g., benzene, toluene, benzoate, and protocatechuate, Supplementary Data 3) were detected in lineages from both terrestrial and aquatic environments (mostly *Houyiibacteriaceae, Anaeromyxobacteraceae*, *Polyangiaceae,* and *Haliangiaceae)*, often coupled with anaerobic respiration using alternative electron acceptors such as nitrate, nitrite, or sulfate (Figure 6, Supplementary Data 4, 5). Additionally, subcellular localization analysis of predicted proteins identified putative polyethylene terephthalate (PET) hydrolases (Supplementary Data 2, 6), further confirmed by NCBI blastp and phylogenetic tree analysis (SI1 Table S4, SI2 Figure S9). While Myxococcota have been recently linked to hydrocarbon degradation in marine and freshwater lineages [5, 34, 111], they are not currently identified as a source of microbial PET hydrolases, highlighting the need for further exploration of their potential in plastic biodegradation. Investigation of the *Houyiibacteriaceae* distribution globally in Sandpiper [47] (v2.1.1 in GTDB r232 taxonomy) revealed the family’s highest relative abundances are in Danish coastal marine sediments (15 of the top 20 metagenomes, and 45 of the top 100, with relative abundance ranges of 5.19 - 18.15%, SI1 Table S20). Their coastal sediment locations indicate likely adaptations to oxygen limited conditions, with exposure to light and pollutants from agriculture and industry. The high abundance of this group in Denmark’s sediments pinpoint this group as an important local target for isolation and application to bioremediation efforts.

## Conclusion

This study proposes that Myxococcota comprise distinct, habitat-structured lineages with fundamentally different ecological strategies. Known predation and multicellular development capabilities, so far considered a hallmark of these microorganisms, are largely observed in soil-associated lineages rather than aquatic lineages, suggesting their evolution as adaption to heterogeneous, nutrient-variable, and less solid surface availability environments. In contrast, aquatic lineages exhibit alternative strategies, including anaerobic metabolism, phototrophy, and expanded roles in biogeochemical cycling. The extensive potential for secondary metabolite production, organohalide transformation, and hydrocarbon degradation highlights a previously underappreciated functional breadth. This is particularly evident in the of genus *Thyboronia* (former g SG8-38) in the family of *Houyiibacteraceae*, with high abundances in Danish coastal environments indicating considerable potential for isolation. Together, these findings position Myxococcota as promising targets for biotechnological applications, emphasizing the need to access the uncultured diversity to unlock their potential in natural product discovery and environmental remediation.

## Author contributions

Conceptualization: F.P., Y.Y., C.M.S., M.A., P.H.N.;

Methodology: F.P., Y.Y., Z. K., C.J., T.B.N.J., M.S., A.D., K.S.K., F.D.;

Investigation: F.P., Y.Y., Z. K., C.J., A.D., K.S.K.;

Supervision: C.M.S., P.H.N.;

Writing of original draft: F. P., Y.Y., Z.K.;

Review and editing: F.P., Y.Y., Z. K., C.J., T.B.N.J., M.S., A.D., K.S.K., F.D., M.A., C.M.S., P.H.N.

## Data availability

The 10,683 MFD short-read Illumina metagenomes and 16S rRNA gene reads were collected from the NCBI GenBank under BioProject PRJNA1071982. The 186 Myxococcota MAGs were collected from the Microflora Danica long-read Nanopore metagenomes dateset at ENA with BioProject ID: PRJEB58634. The 69 Danish MiDAS Illumina metagenomes were collected from the NCBI SRA and GenBank databases under the bioproject accession number PRJNA629478. The 29 recovered Myxococcota representative MAGs in the Danish MiDAS dataset were collected from Figshare under DOI 10.6084/m9.figshare.c.5277035. All data generated during the study are included as supplementary files.

## Code availability

The methods above indicate the source and versions of the programs and code used for the analyses.

## Acknowledgements

We would like to thank the Microflora Danica Consortium for their great help in collecting samples and metadata.

