## Supplementary Information 2 for "From soil to sea: unravelling the metabolic versatility and social dynamics of Myxococcota bacteria from different Danish environments"

#### for

**Running Title:** Myxococcota ecology from soil to sea

**Contents:**

Supplementary Methods

Supplementary Note

Figure S1. Genome sizes of Myxococcota.

Figure S2. Maximum-likelihood (PhyML) 16S rRNA gene phylogenetic tree of Myxococcota species from different Danish environments.

Figure S3. Distribution of Myxoccota across Danish environmental habitats.

Figure S4. Quantification of Myxococcota genomes across Danish habitats.

Figure S5. Distribution of selected Myxococcota families across a broad range of habitats in Sandpiper.

Figure S6. Novelty of Myxococcota.

Figure S7. Number of BGCs detected per HQ Danish MAG.

Figure S8. Phylogenetic tree of DsrAB.

Figure S9. Phylogenetic tree of functionally verified polyesterases having activity on PET.

#### Supplementary Methods

##### *Descriptions of sample metadata, habitat classification, and genome reconstruction*

Dereplicated genomes recovered from 177 sequenced Danish environmental samples were collected from MFD [1] and Danish MiDAS genome [2] projects. Companied with these samples are detailed metadata of the types of sample (i.e., soil or sediment), area (i.e., urban or natural), and a three-level habitat ontology (i.e., MFD ontology level 1/2/3 - MFDO1/2/3 - Bogs, mires and fens - Calcareous fens - Alkaline fens). In the MiDAS genome project, genomes of activated sludge (AS) in 23 Danish wastewater treatment plants (WWTPs) were reconstructed using the hybrid approach with both short-read and long-read sequencing data. In the MFD project, genomes in 154 deep long-read metagenomes (median of 94.9 Gbp sequencing data per sample) were reconstructed using mmlong2-lite [41] (v.1.0.2). Genomes recovered from these projects were originally dereplicated using dRep [3] (v2.3.2 & v2.6.2) at 95% average nucleotide identity genome clustering for distinct species, yielding > 20,000 medium- (MQ) to high-quality (HQ) Danish MAGs following minimum information about a metagenome-assembled genome (MIMAG) criteria [4]. From them, a total of 215 MQ and HQ Myxococcota species representatives were recovered (186 from long-read MFD [1] and 29 species representatives from MiDAS [2]) with taxonomy based on GTDB-tk (v2.4.0) [5] (v2.4.1) using the conserved marker genes defined in GTDB [6] R226 release.

##### *Genome mining of antimicrobials*

The web-based classification tool NaPDoS2 [7] was used with the default cutoffs: blastp, e-value ≥1e-8 and ≥200 aa match length against the database of NaPDoS2_v13b and additional cutoff of ≥50% aa identity.

##### *FISH probe design*

Optimal formamide concentration for each FISH probe was determined after performing hybridization at different formamide concentrations in the range 0-70% (with 5% increments). The intensity of at least 50 cells was measured using ImageJ software [8].

##### *Nomenclature for naming*

Generally, names of the Danish sampling locations/cities were used to generate the latinised genus names, and then propagated to the higher taxonomic level naming whenever the upper taxonomic levels have placeholder names. For the novel class UBA9160, we named it after Martin Vahl from the Flora Danica project that inspired the initiation of the MFD project [1,9]. The incorrect spelling of the class Polyangia has been fixed to Polyangiia [10].

### Supplementary Note

**Potential for predation machineries in Myxococcota from diverse environments.**

**Motility and chemosensory systems**. Myxococcota's predatory lifestyle begins with a remarkable chemotactic ability that enables individual cells to sense and respond to environmental cues. As nutrient levels fluctuate, these bacteria initiate coordinated movement, forming swarms that navigate their surroundings in search of prey and lyse them, while the predator remains unharmed. The organic matter released from the preys is then degraded, taken up, and metabolized [11,12]. However, despite being for a long time considered as a typical trait of the phylum, predation and social behavior may not be as diffused as originally thought. A recent study predicted based on genomic evidence that uncultured Myxococcota from non-soil habitat may display a diverse metabolic flexibility, while the predatory machinery may be a consequence of the adaptation to terrestrial ecosystems[13]. We assessed the distribution patterns of pathways involved in predation, including motility, chemosensory systems, secretion systems, production of extracellular exopolysaccharides (ExoPS), hydrolytic enzymes and secondary metabolites.

Motility is a crucial mechanism in the predatory lifestyle of Myxococcota and is generally characterized by two different movement strategies, namely adventurous (A-motility) and social motility (S-motility). In the first, *M. xanthus* single cells glide on solid surfaces powered by proton motive force (PMF) and a multiprotein complex (AglRQS) that span cytoplasm, membrane and periplasm[14–16]. The flagella motor complex AglRQS is formed by AglR, homolog of *E. coli* MotA, and two MotB homologs, AglQ and AglS, and is linked to cytoplasmic filaments (MreB). The motor complexes are hypothesized to carry protein cargos (AgmU and AglZ) along these filaments and exert force against the cell envelope through these, without breaching through the peptidoglycan layer[15,16]. Homologs of MotA, MotB and the other accessory proteins were widespread in the MAGs associated to the *Myxococcia* class, with the only exception of some MAGs belonging to the *Anaeromyxobacteraceae* and 40CM-4-68-19 families, as well as in Kuafuiibacteriia and the novel class UBA727 (Figure 3). The potential for A-motility did not appear to be ecosystem-related, as it was present not only in families predominant in soils, such as *Myxococcaceae* and *Blaeremorphaceae*, but also in several MAGs retrieved from freshwater environments (Figure 3). Several MAGs possessed only homologs of the subunit MotA or the motility proteins were absent (Supplementary Data 1), indicating potential absence of the A-motility, as observed in some isolates that only exhibit S-motility[17].

In *M. xanthus* S-motility, polymerized type IV pili are extended at one pole of the cell to act as “hooks" and then retracted to pull the cell forward, typically in a self-secreted matrix of exopolysaccharides. Type IV pili machines consist of more than 10 conserved core proteins (Pil) spanning from the cytoplasm to the outer membrane[16,18,19]. Homologs of the Pil machineries were present in all the MAGs, with few exceptions in the families *Houyiibacteriaceae*, *Sandaracinaceae*, *Blaeremorphaceae*, and *Nannocystaceae* (Figure 3). However, the prediction of type IV pilus assembly is not an unambiguous indication for S-motility potential, as these cell structures may be involved in other processes, such as adhesion, secretion and uptake, or virulence[14,20]. S-motility in *M. xanthus* is also adjuvated by ExoPS secretion[16]. Homologs of the *eps* gene cluster from *Pseudomonas* were detected in several MAGs of the class Myxococcia*,* as well as some representatives of the class Polyangiia (family *Polyangiaceae*) (Figure 3). S-motility is also ensured by a specialized chemosensory network, which coordinates and synchronizes cell movements and consists of the proteins Dif, Frz, Mgl and RomR[13,21]. Genes encoding for homologs of Dif and Frz were widespread in the MAGs belonging to the class Myxococcia*,* but only sporadically detected in the other classes (Figure 3), and only the GTPase MglAB and the response regulator RomR were widespread throughout the phylum (Supplementary Data 1). No homologs for the Frz module were identified in *Haliangium ochraceum* and other *Haliangiaceae* MAGs, despite the experimental evidence for social motility in the isolate. These results could indicate that other Myxococcota with social motility may use a different type of cell-to-cell interaction to synchronize their motility, likely involving only homologs of the chemotaxis system Dif, which is more widespread throughout the phylum (Figure 3).

Cell-to-cell interactions are also enabled by outer membrane exchange (OME) mechanisms, which allows resource sharing as well as self/non-self discrimination[22,23]. In *M. xanthus* cells, self-recognition is mediated by contact between two identical outer membrane receptors TraA and the accessory protein TraB. The transient membrane fusion of the TraA receptors leads to the bidirectional transfer of large amounts of outer membrane proteins and lipids, as well as diverse SitA toxins. The *sitA* genes in the genome are generally accompanied by a downstream immunity gene (*sitI*), which are expressed together and ensure immunity for *M. xanthus* cells, but are not transferred during OME, ensuring that OME can happen only between clonal cells [13,22,23]. While homologs of TraB were widespread throughout the phylum, TraA was only presents in few MAGs of the class Myxococcia (*Myxococcaceae,* f__UBA5297, o__SLRQ01), c__UBA727, Kuafuiibacteriia,, c__B64-G9, Bradymonadia and Polyangiia (*Nannocystaceae*, *Haliangiaceae*, *Sandaracinaceae* and *Polyangiaceae*) (Figure 3). SitA genes were mainly observed in *Myxococcus* close relatives, indicating that the production and exchange of this toxin may be limited to this lineage (Supplementary Data 1).

**Hydrolytic enzymes and secretion.** *M. xanthus* uses a combination of secreted enzymes and secondary metabolites to kill and lyse prey cells[24–26]. The genome encodes potential for peptidoglycan degradation, which could serve as killing factors that induce prey cell lysis, and proteolytic activity, to degrade proteins and peptides released from prey cells. Amidases and glucosaminidases have been isolated from *M. xanthus* cultures supernatant early on[11,12], and an outer membrane β-1,6-glucanase from the myxobacterium *Corallococcus* was shown to be required for lysing the chitinous cell wall of certain fungi[11,18,27]. To explore the predicted extracellular proteins, we performed subcellular location analysis of the encoded proteins using PSORTb [28]. The presence of predicted extracellular peptidases/proteases was widespread throughout the phylum, with few exceptions in the class Polyangiia (*Polyangiaceae*, *Houyiibacteriaceae*, *Blaeremorphaceae,* and f__Palsa-1140), in the family *Anaeromyxobacteraceae* and in the classes c__UBA727 and Vahlivitia (Figure 3). The predicted extracellular peptidases belonged mainly to the MEROPS peptidase families M23, M28 and M4, and serine peptidases S8 and S1 (Supplementary Data 5). These gene families include peptidase enzymes that can cleave the peptide bonds in the peptidoglycan of bacterial cell walls, as well as bacteriocins and thermolysin-like peptidases, known for their extracellular protein digestion function or for their effects as inhibitors of protein synthesis and antibacterial activity against other bacteria [13]. Extracellular carbohydrate active enzymes (CAZy) were also predicted in the majority of the MAGs, with a prevalence of glycoside hydrolases (GH) of the families GH5 and GH9, which include endoglucanases and cellobiohydrolases for cellulose degradation, and polysaccharide lyases (PL) for pectin degradation (Supplementary Data 1). While model Myxococcota genomes are generally enriched in GH23 and GH13 (peptidoglycan lyases and amylases), our results indicate that extracellular CAZymes predicted in our MAGs are likely involved in digesting the diverse extracellular polysaccharides, rather than in directly killing prey cells. Predicted extracellular lipases and nucleases were less common and, when present, likely involved in digesting the cell wall lipids and intracellular content of their prey (Figure 3, Supplementary Data 1).

Targeted secretion via protein secretion systems of the types II, III, IV, or VI are generally used in other predator bacteria to deliver hydrolytic enzymes in the proximity or directly into prokaryotic or eukaryotic cells[20,27]. Our MAGs carried a wide range of secretion systems, with several MAGs encoding for the apparatus proteins of T2SS, with the exceptions of few MAGs in the *Nannocystaceae* and *Anaeromyxoobacteraceae* families. *M. xanthus* utilizes T3SS, T3SS(2) and a Tad-like secretion complex to export digestive enzymes from the cells[20,29,30]. However, homologs of these secretion systems were mainly detected in the MAGs belonging to the class Myxococcia. T6SS and Tad-like secretion machineries appeared to be more widespread across the phylum (Figure 3, Supplementary Data 1). While experimental evidence showed that mutations of the T6SS did not affect the predation abilities in *M. xanthus*[20], they could be involved in cell contact-dependent predatory functions in lineages outside of the *Myxoccocaceae* family.

**The biosynthetic antimicrobial potential.** Model Myxococcota organisms also secrete a plethora of secondary metabolites, such as pigments, siderophores, bacteriocins, and antimicrobials, that attack and lyse their prey. AntiSMASH detected a total of 918 BGCs from these 90 HQ Danish Myxococcota MAGs (SI1 Table S11), the majority of which were complete (821/918, 89.4%). The highest counts of BGCs per MAG were found in the families *Sandaracinaceae*, CAJQPC01, *Nannocystaceae*, and VGSZ01 (Figure 1, SI2 Figure S7). At the individual MAG level, the highest counts of BGCs per MAG were observed in the families of *Haliangiaceae* (27 BGCs), *Myxococcaceae* (23 BGCs), and *Sandaracinaceae* (22 BGCs). Notably, only one quarter of these BGCs had a match to known BGCs in the MiBIG database (225/918). Furthermore, only 27.6% (62/225) of the BGCs had a similarity precision proportion >50% to a known BGC, adding to the high novelty of Danish Myxococcota BGCs. The detected known BGCs spanned the biosynthesis of 80 compounds (SI1 Table S12), with most related to pigments (e.g., 1 isorenieratene, 32 carotenoids, 7 aryl polyenes, 1 zeaxanthin), fruiting body formation (i.e., 31 VEPE/AEPE/TG-1), exoPS formation (e.g., 5 N-tetradecanoyl tyrosine, 3 lipopolysaccharide, 5 heteropolysaccharide, 2 exopolysaccharide), survival under stress (e.g., 6 ectoine, 5 hopene), antimicrobials (e.g., 3 microsclerodermin, 1 lankacidin, 2 asukamycin, 2 bottromycin, 2 crocacin, 1 stigmatellin, 1 leinamycin). Among the known BGCs, 12.4% (28/225) were linked to antimicrobial biosynthesis. These BGCs represented a third of the currently known antimicrobial compounds, reflecting the potential of environmental Myxococcota for new antibiotic compounds discovery [31].

We further examined the biosynthesis potential for antimicrobial compounds by identifying the highly conserved and functionality-informative C domains in NRPS, and KS domains in PKS using NaPDoS2 (Figure 4) for all AntiSMASH-identified BGCs. A total of 233 KS and C domain sequences (with min. 50% aa identity) were detected in 81/90 of the Danish Myxococcota (SI1 Table S13). These domain sequences spanned the biosynthesis of 47 compounds, within which half of the compounds were reported with antimicrobial property (26/47, Figure 4) found in a fifth of the domain sequences (55/233) located on 21/90 HQ Danish Myxococcota. The most detected domain sequences encoded antimicrobial compounds were for stigmatellin (7/55), chondrochloren (6/55), leinamycin (6/55), and sanglifehrin (5/55), inhibiting a broad range of both Gram-positive and Gram-negative bacteria (Figure 4). Overall, homology (53-67% aa identity, $10^{th}$-$90^{th}$ percentile) of all the 55 identified domains to characterised antimicrobial domains supports the potential of Myxococcota to use antimicrobials as a predatory machinery.

**Cellular differentiation and sporulation mechanisms in Myxococcota from different environments.**

Many Myxococcota display a facultative multicellular behaviour in response to environmental cues. When the nutrients are insufficient, Myxococcota cells move in swarms using S-motility and aggregate into mounds, with cellular differentiation to spores in fruiting bodies[32–34]. These resistant spores are, however, ready to sporulate in response to nutrient signals[32]. In *M. xanthus*, this complex multicellular developmental process is regulated by a sequence of signal-responsive transcription factors involved in the aggregation, mold formation and sporulation[21,33,35,36]. Starvation is activating a cascade of phosphorylation of several transcription factors (in *M. xanthus* Nla4, Nla18, Nla6, and Nla28), which themselves activate the Act and FruA modules[13,21,35,36]. Homologues for these transcription factors were identified in several of our MAGs (Figure 3, Supplementary Data 1), however this may likely be due to the common presence of DNA-binding transcription factors and protein kinases domains in bacterial genomes [13]. The diguanylate cyclase dxmB the transcription factor Nla24 are also activated by starvation and induce activation of the *eps* gene cluster and production of ExoPS, necessary for aggregation and fruiting bodies formation[13,21,32,35,36]. Homologues for dxmB and Nla24 were widespread throughout the phylum (Figure 3, Supplementary Data 1), but occurrence of diguanylate cyclase and DNA-binding transcriptions factors domains are also very common in bacterial genomes [13]. The presence of ExoPS production potential (*eps*), together with the transcription factors involved in these early stages of aggregation may be a more precise indication of fruiting bodies formation. Finally, the Mrp network (MrpA, B, and C) is also activated in response to starvation and activates and acts cooperatively with the FruA module, controlling the expression of several genes essential for aggregation and/ or sporulation[13,21,35,37]. Homologs for these genes were identified in several MAGs of the *Myxococcaceae*, *Haliangiaceae* and *Nannocystaceae* families (Figure 3, Supplementary Data 1). Some MAGs encoded genes for MrpA and/or MrpB, but not the main transcription factors MrpC or FruA, indicating an unlikely probability of aggregation/sporulation in these lineages (Supplementary Data 1). Since these proteins are histidine kinases or DNA-binding transcriptional response regulators, they may likely be involved in other processes[13].

Several operons are involved in the sporulation, including the Fmg, Exo, mcu, Nfs operons and the spore-coat protein Tps, involved in forming the spore polysaccharide coat, as well as several signal transduction systems (RodK/RokA, RedCDEF), coordinate the timing of aggregation and sporulation[13]. Only MAGs belonging to the class Myxococcia encoded the full operons controlling sporulation, but some sporulation genes homologues were identified in other MAGs (e.g. families *Polyangiaceae*, *Nannocystaceae*, *Haliangiaceae,* and Kuafuiibacteriia). Several isolates in these families are known to be spore-formers and have fruiting bodies[38–40], such as *H. ochraceum* and *N. exedens*, but lack homolog genes for the aggregation and sporulation processes, indicating a potential different mechanism at molecular level. Therefore, we cannot exclude that other genera outside of the family *Myxococcaceae* could exhibit some forms of multicellular development.

**Myxococcota possess a versatile lifestyle in different environments**

Functional analysis of the 238 (215 Danish MAGs and 23 GTDB isolate MAGs) Myxococcota genomes revealed a versatile metabolism, reflecting their adaptation capabilities and likely their survival, and sometimes predominance, in both eutrophic and more extreme environments. Generally, cultured members of the Myxococcota phylum possess a heterotrophic lifestyle. Our results confirmed this feature, as the genomes encoded complete or almost complete central carbon processing, through glycolysis, pentose phosphate pathway and TCA cycle (Figure 6, Supplementary Data 2). Potential carbon sources included different carbohydrates, with potential for uptake of glucose/mannose (*msmX*), trehalose/maltose (*thu*), and in fewer MAGs also fructose (*frc*) and xylose (*xyl*) (Supplementary Data 2). Putative multiple sugars transporters were also widespread throughout the phylum (Supplementary Data 2). Their potential for carbohydrate degradation is also confirmed by the possession of multiple glycoside hydrolases and polysaccharide-degrading CAZymes (Supplementary Data 2). Most Myxococcota isolates rely mainly on amino acids and lipids as carbon sources[12,13,41–43]. Our results confirmed that peptides and amino acids are essential substrates, with detection of multiple copies of peptides and oligopeptides transport systems (*opp*), branched amino acids (*liv*) and glutamate transporters (*glt*), as well as several amino acids degradation pathways (Figure 6, Supplementary Data 3). Several MAGs across the phylum exhibited the ability to uptake and consume acetate, through the phosphate acetyltransferase-acetate kinase pathway (*actP* for uptake, *acs*, *pta* and *ackA* for utilization) (Supplementary Data 3). Similarly, the potential for a complete glyoxylate cycle was also widespread (Supplementary Data 2), and several MAGs also encoded the full potential for aerobic pyruvate oxidation to acetyl-CoA (*aceE* and *aceF*). Lipids are also a common substrate for model Myxococcota[13,37,44], but none of our MAGs possessed a complete beta-oxidation pathway for long-chain fatty acid degradation, suggesting that these compounds may not be one of their preferred energy sources (Supplementary Data 2).

As several Myxococcota families have been recently identified as phototrophs [45], we investigated potential for carbon fixation in our MAGs. Some members of the *Myxococcaceae*,

*Anaeromyxobacteraceae*, *Polyangiaceae*, *Nannocystaceae*, f__UBA5297, f__Palsa-1140 and *Blaeremorphaceae* possessed the potential for a complete or near complete Calvin-Benson-Bassham (CBB) cycle (Figure 6), characterized by a type I ribulose-1,5-bisphosphate carboxylase/oxygenase (RuBisCO), generally found in other photosynthetic bacteria [46]. We identified full bacteriochlorophyll biosynthesis and light harvesting pathways in five MAGs (one in *Kuafuiibacteriaceae*, one in f__Nannocystaceae, two in f__SG8-38 and one in f__40CM-4-68-19) retrieved from activated sludge, marine sediments or bogs soils (Figure 6, SI1 Table S15). Phototrophy may play a key role in helping these Myxococcota survive when prey or organic nutrients are scarce and incorporating phototrophic traits may help these bacteria meet energy demands for movement, enhance secondary metabolite production, and better adapt to environmental stresses through a flexible light-based lifestyle. These results confirm previous findings on potential photosynthetic activity of these microorganisms and it is further increasing the number of families with phototrophic capabilities in the Myxococcota [46,47]. Several biosynthetic gene clusters for production of carotenoids and other pigments were detected in our MAGs (Figure 6, SI1 Table S16), and Myxococcota isolates are generally known for their peculiar pigmentation[24,45]. Biosynthesis of beta-carotene, zeaxanthin and isorenieratene were identified in about a third of Danish Myxococcota genomes (77/215), spanning a wide range of Myxococcota classes (i.e., c__Myxococcia, c__Polyangiia, c__Bradymonadia, and Vahlivitia) and a broad range of habitats, including activated sludge, harbour saltwater, and various types of soils and sediments (SI2 Figure S3), likely serving in phototrophic process as accessory pigments and protection against oxidative stress, but also as antimicrobial active compounds[13,45].

The majority of the cultured Myxococcota, with the exception of *Anaeromyxobacter*, are aerobes. However, functional analysis of our MAGs showed potential for facultative and, in few cases, anaerobic lifestyle. Potential for fermentative processes of pyruvate were present in several MAGs, with ethanol or lactate as by-products (Figure 6, Supplementary Data 3). Few MAGs also encoded the genes for a formate dehydrogenase (*fdh*), potentially used to reduce formate produced during anaerobic fermentation (Supplementary Data 3). Almost all our MAGs encoded the homolog genes for a cytochrome c oxidase (*cox* and/or *coo*), with few exceptions in the class Vahlivitia, and the families *Houyiibacteriaceae*, *Myxococcaceae*, *Polyangiaceae*, *Anaeromyxobacteraceae*, *Vahlivitaceae* and *Blaeremorphaceae* (Figure 6, Supplementary Data 3). Cytochrome bd ubiquinol oxidase was often encoded in the MAGs lacking other cytochrome c oxidases (Supplementary Date 3), but it is proposed to have a detoxification role in strictly anaerobes[13]. The Wood Ljungdahl pathway (WLP) has been previously detected in Myxococcota MAGs and proposed as an electron sink mechanism to re-oxidize reduced ferredoxin and produce additional ATP[13], but none of our MAGs encoded a full WLP. However, some of the MAGs encoded the *Rhodobacter* nitrogen fixation (RNF) complex genes and the cytoplasmic electron bifurcating mechanism HydABC, which can potentially be used to generate proton motive force and ATP in anaerobic bacteria[13].

**Myxococcota possess a potential ecological role in bioremediation processes**

Microorganisms´ metabolic abilities, including the breakdown of pollutants and the production of secondary metabolites, have driven advancements in biosynthesis and biodegradation, providing solutions to various environmental issues. Uncharacterised microorganisms possess a vast repository of undiscovered enzymes that hold the potential for applications in various fields, including biotechnology and environmental remediation. Myxococcota are known to possess a variety of diverse metabolic abilities, including nitrogen cycling[17,48], aromatic hydrocarbon degradation[49], hydrogenotrophic respiration[49] and organohalide respiration[17].

Our MAGs encoded the genes for potential involvement in the nitrogen cycling. Alternative terminal electron acceptors under anoxic conditions included nitrate and nitrite, through dissimilatory nitrate reduction (with either NarGHI or NapAB enzymes) and/or nitrite reduction via the NirK/NirS enzymes (Figure 6, Supplementary Date 3). Few MAGs also encoded potential for nitric oxide reduction to nitrous oxide (NorBC) and for potential reduction of the latter to gaseous nitrogen (NosZ) (Figure 6, Supplementary Date 3). Only few MAGs possessed the full potential for denitrification, one from saltwater sediments (Vahlivitia), one from freshwater sediments (genus *Anaeromyxobacter*), one from natural soil (*Polyangiaceae*) and four from activated sludge (f__Ga0077539, *Polyangiaceae* and *Haliangiaceae*) (Figure 6, Supplementary Date 3). Additionally, some of the MAGs encoded the potential for dissimilatory nitrite reduction to ammonia via the cytochrome-linked nitrite reductase NrfAH (Figure 6, Supplementary Data 3). Only a few genomes (32/215) encoded the capability to fix atmospheric nitrogen (Figure 6, Supplementary Data 3). Within Myxococcota, nitrogen cycling has largely been studied in *Anaeromyxobacter dehalogenans*[17,48,50] and has been suggested for *Haliangium*-related species abundant in activated sludge[51], but these pathways are not well characterized in other lineages within this phylum. Phosphorus is also an essential element for life and Myxococcota have a potential role in phosphorus cycling, through the uptake of inorganic P from the environment and synthesis of intracellular polyphosphate (polyP). *Myxococcus xanthus* produces polyP through the activity of polyphosphate kinase 1 (Ppk1) and breaks down short- and long-chain polyP using the exopolyphosphatases Ppx1 and Ppx2, respectively [52–54]. Additionally, the polyP:AMP phosphotransferase (Pap) enzyme in *M. xanthus* catalyzes the formation of ADP from AMP and polyP[52,54]. In this organism, polyP plays a significant role in supplying energy for motility, predation and maturation of spores[54].

The genes essential for phosphate uptake (Pit and Pst transporters) and polyP functions were detected in several MAGs from the family *Myxococcaceae*, *Haliangiaceae*, *Polyangiaceae,* phylum Myxococcota_A, and many more (Figure 6, Supplementary Data 3), and prevalent in MAGs from aquatic environments. Progress in understanding the phosphorus biogeochemistry of changing aquatic ecosystems are needed to better comprehend, predict, and address the consequences of human influence[55].

Recent metagenomic studies have revealed that some members of the phylum Myxococcota harbor genetic potential for diverse sulfur metabolic processes, suggesting that they may play important roles in sulfur cycling across a range of environments[13,44]. While assimilatory sulfate reduction was common, only few MAGs (*Blaeremorphaceae*, f__UBA5297 and Vahlivitia) encoded a complete dissimilatory sulfate reduction pathway (Figure 6). In addition, several MAGs harbored genes for tetrathionate and thiosulfate reduction, indicating the capacity to use these reduced sulfur species as terminal electron acceptors under anoxic conditions (Figure 6, Supplementary Data 3). Few Myxococcota MAGs (from *Vahlivitaceae* and *Hyalangium*) also encode the Sox (sulfur oxidation) enzyme complex (Figure 6, Supplementary Data 3), which could facilitate both mixotrophic growth in oxygen-limited niches and contribute to the regeneration of sulfate for other microorganisms, reinforcing their ecological role in coupled carbon–sulfur [56].

The widespread contamination of groundwater and soil with toxic and carcinogenic halogenated compounds poses a significant environmental and public health challenge. Approaches such as biostimulation and bioaugmentation are nowadays widely applied to accelerate the cleanup of contaminated sites, highlighting the valuable role of microbes in sustainable remediation technologies [57]. Organohalide-respiring bacteria, such as *Dehalococcoides* spp., *Dehalobacter* spp., and *Dehalogenimonas* spp., play a key role in driving various dehalogenation reactions [57]. They gain energy for growth through a strictly anaerobic process where they generally utilize hydrogen or formate as electron donors, acetate as a carbon source, and organohalides as electron acceptors [58]. Their genomes can carry more than 30 distinct genes coding for reductive dehalogenases [58], potentially active in the dechlorination of various chlorinated compounds. Within the Myxococcota phylum, reductive dehalogenase encoding genes have been identified in *A. dehalogenans*, which is capable of reductive dechlorination of a range of halogenated aromatic compounds, contributing to the detoxification of chlorinated pollutants in soil and sediment environments [59]. Several homologs of the reductive dehalogenase protein complexes included in the RDaseDB [60] were identified in 37 Danish MAGs, with the majority in the families *Anaeromyxobacteraceae*, *Myxococcaceae*, *Polyangiaceae*, *Houyibacteriaceae,*

*Blaeremorphaceae* and class Vahlivitia (Figure 6, SI1 Table S17). In addition to respiratory dechlorination, various non-respiratory bacterial enzymes, such as haloalkane dehalogenases or 2-haloacid dehalogenases, also remove chlorine atoms from organic molecules [61]. Unlike reductive processes, these reactions do not yield energy for the organism, except for releasing organic carbon that can be further metabolized [61]. Multiple copies of genes coding for potential hydrolytic or oxidative dehalogenases were annotated in the same MAGs (Figure 6, SI2 Table S18). In bioremediation, efforts typically focus on stimulating respiratory reductive dehalogenase genes; however, the frequent occurrence of non-respiratory hydrolytic and/or oxidative dehalogenase genes suggests that they may also play a significant role in the global chlorine cycle and could be valuable for bioremediation, particularly at low organochlorine concentrations [61].

Myxococcota potential involvement in degradation of hydrocarbons has been recently observed in marine and freshwater lineages [13,62,63]. Our genomic analysis revealed genes encoding enzymes for the transformation of hydrocarbons, with complete or nearly complete pathways for benzene (f__UBA5297), benzoate (*Houybacteraceae)*, toluene (mostly in *Anaeromyxobacteraceae*, *Polyangiaceae* and *Haliangiaceae*) and protocatechuate degradation (*Anaeromyxobacteraceae)* (Figure 6, Supplementary Data 2). In some of these lineages, hydrocarbon degradation appear to be coupled with anaerobic or facultative respiratory processes, using alternative electron acceptors like nitrate, nitrite or sulfate, while others may integrate phototrophic energy acquisition or mixotrophic growth to supplement energy demands under nutrient-limited conditions. This metabolic flexibility enhances their resilience in complex microbial communities and supports potential synergistic interactions with other microorganisms that depend on the by-products of hydrocarbon degradation.

Subcellular localization analysis of predicted proteins in the MAGs revealed occurrence of putative polyethylene terephthalate (PET) hydrolases (Supplementary Data 5), suggesting that members of this phylum may contribute to the biodegradation of synthetic plastics in soil and aquatic environments. Currently, Myxococcota are not identified as a source of microbial PET hydrolases, but considering the biotechnological importance of this process, further research is needed to explore the potential of Myxococcota in PET degradation.

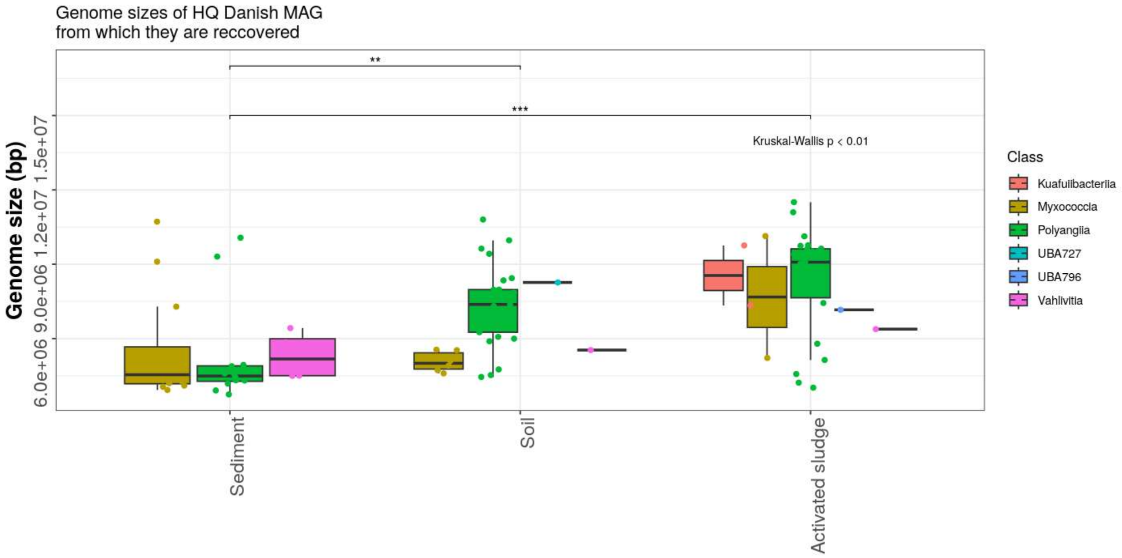


**Figure S1. Genome sizes of Myxococcota.** Boxplots display the distribution of genome sizes for Danish HQ MAGs recovered from different sample types, stratified to different classes. Kruskal-Wallis Rank sum test shows significant difference in genome sizes among these 3 different sample types, and whiskers indicate the significant difference between sample types (**p <0.01 and ***p < 0.001). Classes UBA796 and Vahlivitia are from phylum Myxococcota_A, and all the rest classes are from phylum Myxococcota.

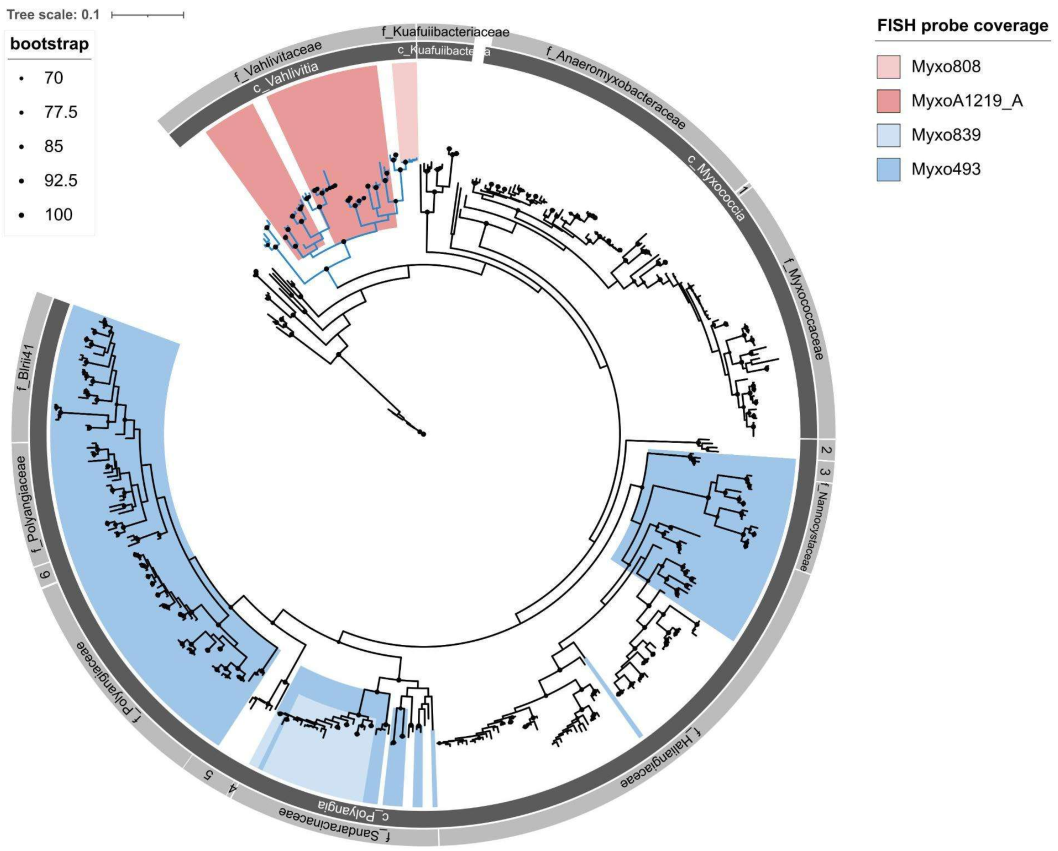


**Figure S2. Maximum-likelihood (PhyML) 16S rRNA gene phylogenetic tree of Myxococcota species from different Danish environments.** 16S rRNA gene sequences used in the tree were retrieved from the genome catalogues of MFD and the MiDAS genome projects. The alignment used for the tree applied a 20% conservational filter to remove hypervariable positions, giving 1159 aligned positions. The blue tree branches (on the top left corner of the tree) represent Myxoccocata_A clade. Taxonomic classification on class and family levels are added in the outer layer (1 - f_Vulgatibacteriaceae; 2 - f_MFD_f_9449; 3 - f_MFD_f_523; 4 - f_MFD_f_18916; 5 - f_MFD_f_1090; 6 - f_Phaselicystidaceae). Coverage of FISH probes is indicated with colored boxes and is based on the MFD reference database (SI1 Table S6). Bootstrap values from 1,000 re-samplings are indicated for branches with >70% support. Species of the phylum Bdellovibrionota were used as the outgroup. The scale bar represents substitutions per nucleotide base. Vahlivitia: prev. Vahlivitia; Blaeremorphales: prev. o__Fen-1088; *Vahlivitaceae*: prev. f__UBA9160; *Houyiibacteriaceae*: prev. SG8-38; *Blaeremorphaceae*: prev. f__Fen-1088; Kuafuiibacteriia; prev. c__WYAZ01; *Kuafuiibacteriaceae*; prev. f__WYAZ01. Classes UBA796 and Vahlivitia are from phylum Myxococcota_A, and all the rest classes are from phylum Myxococcota.

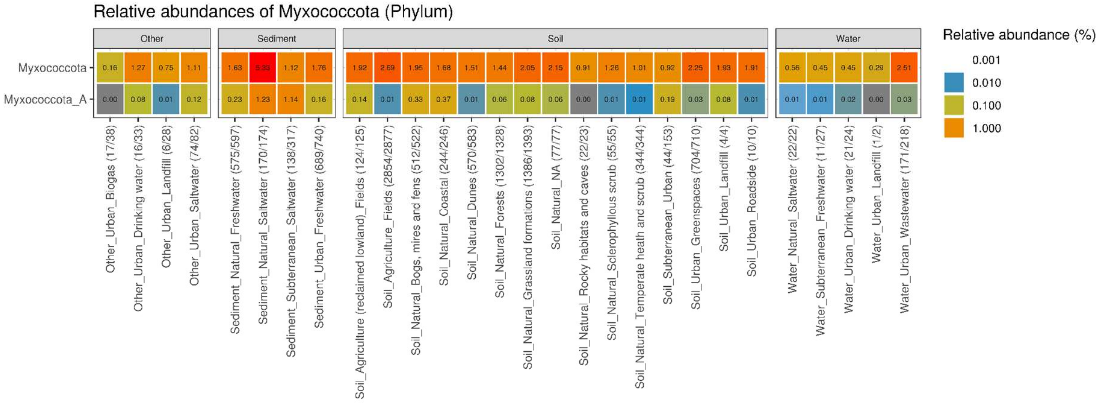


**Figure S3. Distribution of Myxoccota across Danish environmental habitats.** Heatmap showing the average relative abundances of Myxococcota phyla detected in 10163/10752 samples across 28 different Danish habitats (habitat label showing the sample type, area type, and MFDO1). Proportion of samples detected with Myxococcota in different habitats are indicated in the brackets in the x axis.

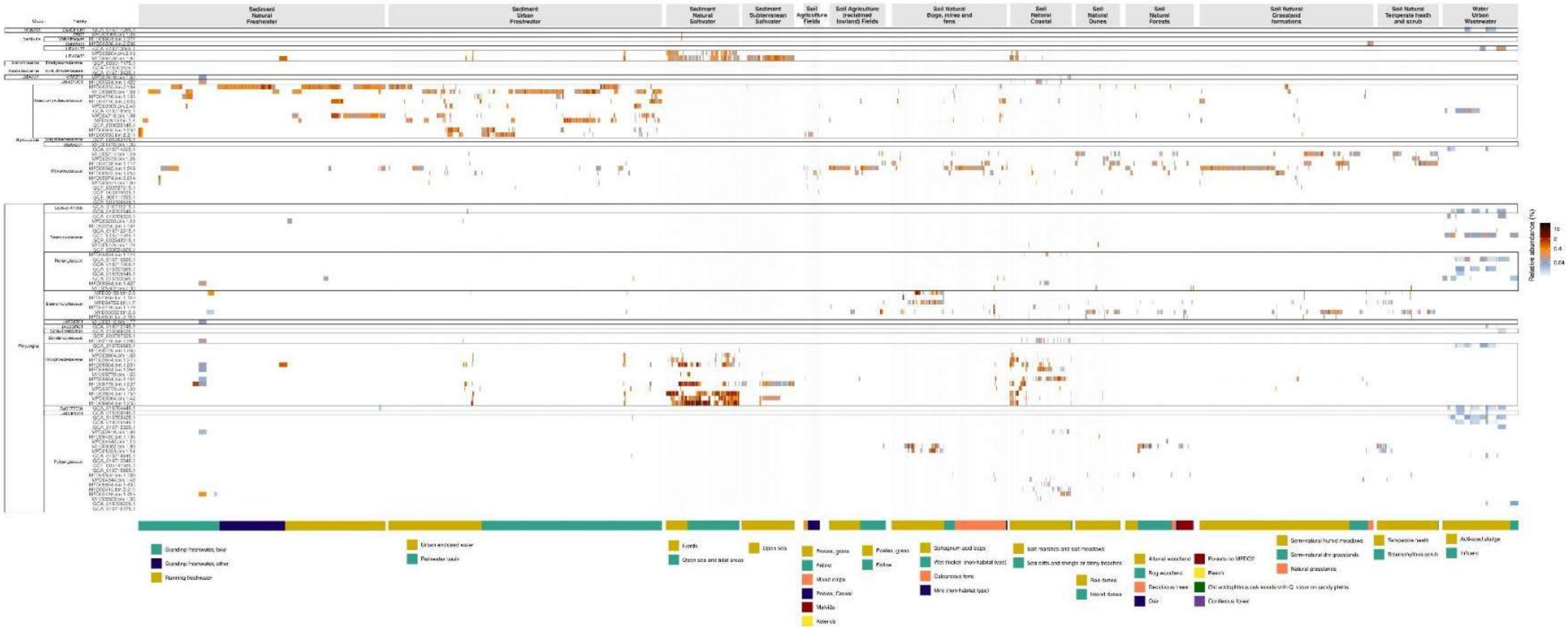


**Figure S4. Quantification of Myxococcota genomes across Danish habitats.** Phylogenomic tree of subsetted Myxococcota genomes from 11 HQ isolates in GTDB and 90 HQ Denmark MAGs. Genomes are separated and taxonomically labelled at the family level on the left. Each genome is quantified by sylph. Only those MFDO1 with at least 10 samples detected with one of these genomes are included. Within each MFDO1, habitats are clustered by MFDO2. Vahlivitia: prev. Vahlivitia; Blaeremorphales: prev. o__Fen-1088; *Vahlivitaceae*: prev. f__UBA9160; *Houyiibacteriaceae*: prev. SG8-38; *Blaeremorphaceae*: prev. f__Fen-1088. Classes UBA796 and Vahlivitia are from phylum Myxococcota_A, and all the rest classes are from phylum Myxococcota. (High resolution version submitted as FigureS4.pdf as supplementary file).

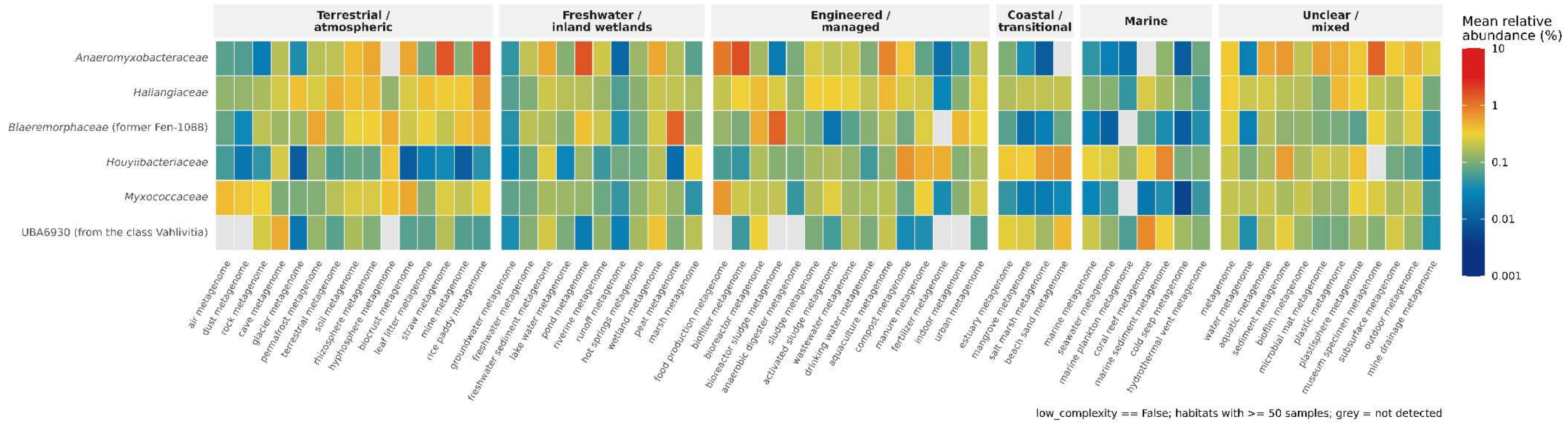


**Figure S5. Distribution of selected Myxococcota families across a broad range of habitats in Sandpiper.** Heatmap showing the mean relative abundances of 6 different Myxococcota families across a varied habitats.

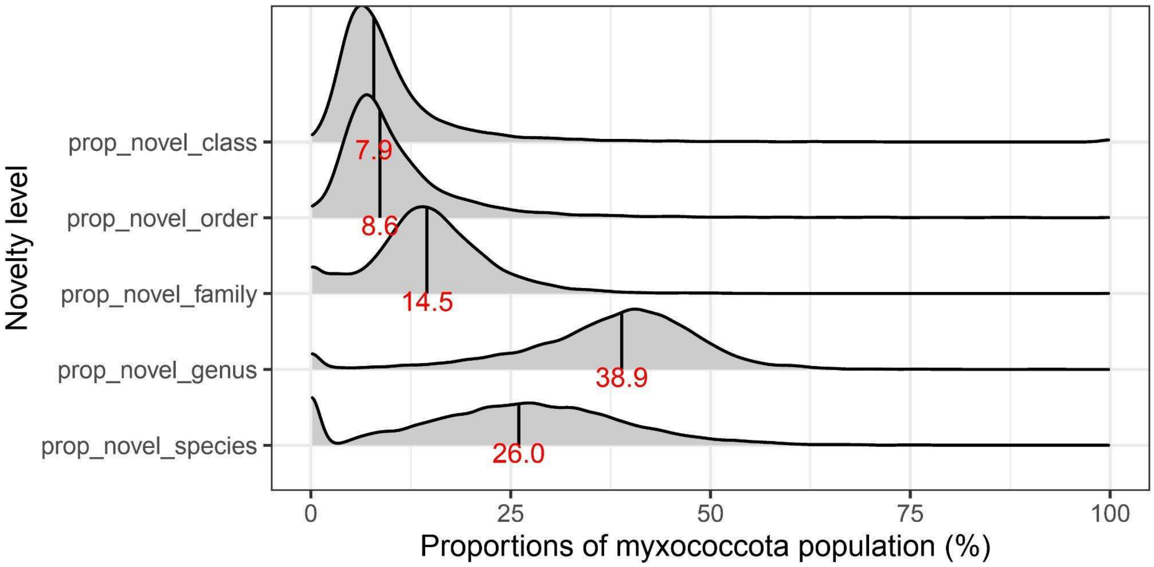


**Figure S6. Novelty of Myxococcota.** Ridgeline plots showing the distribution of the proportion of novel Myxococcota among the Myxococcota population at different taxonomic levels (exclusive: results at different taxonomic levels are not cumulative). Averages are plotted with vertical lines in the ridgeline with values indicated in red.

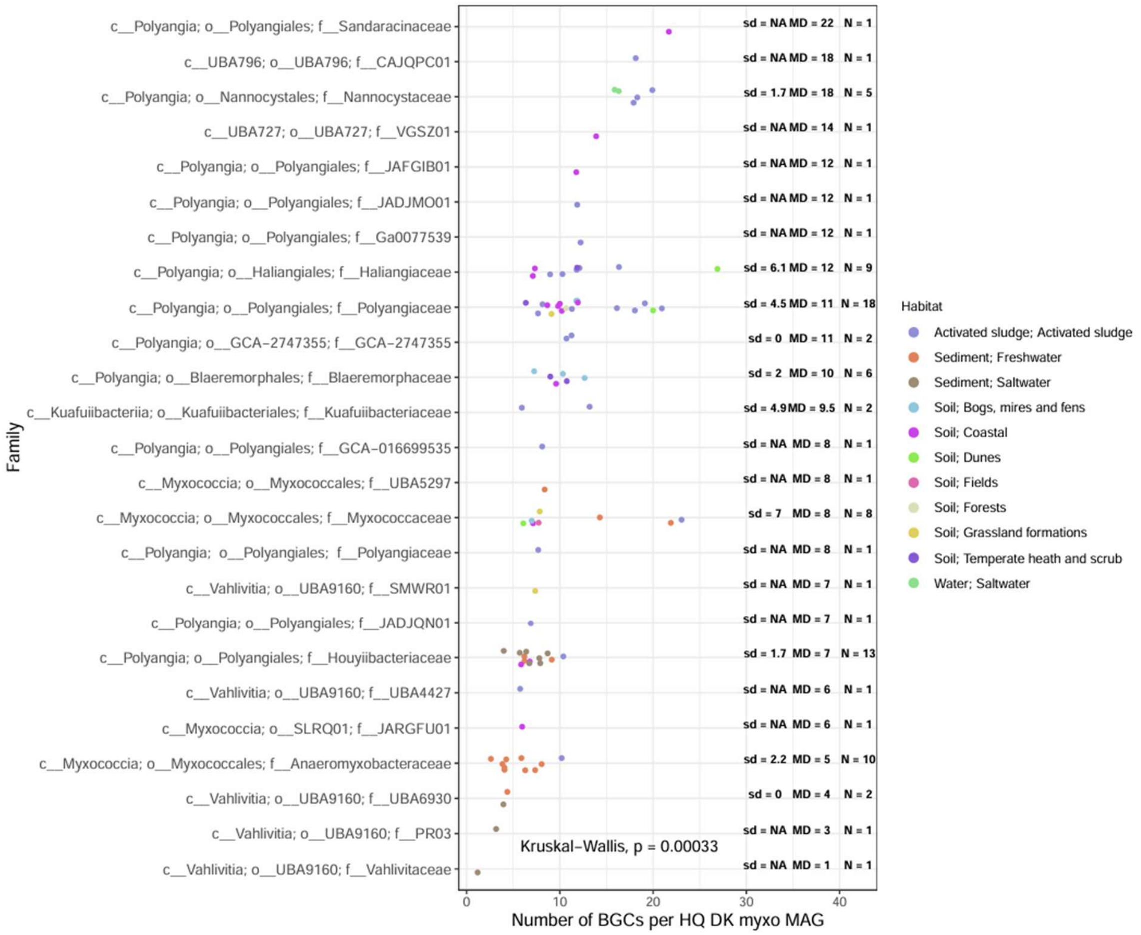


**Figure S7. Number of BGCs detected per HQ Danish MAG**, stratified to different Myxococcota families. MD: median, sd: standard deviation, N: number of MAGs. Kruskal-Wallis test showed that the number of BGCs possessed by different Myxococcota families were significantly different. Habitats indicate where the MAGs were retrieved at the sample type and MFDO1 levels. Vahlivitia: prev. c__UBA9160; Kuafuiibacteriia; prev. c__WYAZ01; *Blaeremorphales*: prev. o__Fen-1088; *Kuafuiibacteriales:* prev. o__WYAZ01; *Kuafuiibacteriaceae*; prev. f__WYAZ01; *Vahlivitaceae*: prev. f__UBA9160; *Houyiibacteriaceae*: prev. SG8-38; *Blaeremorphaceae*: prev. f__Fen-1088 DK: Denmark.Classes UBA796 and Vahlivitia are from phylum Myxococcota_A, and all the rest classes are from phylum Myxococcota.

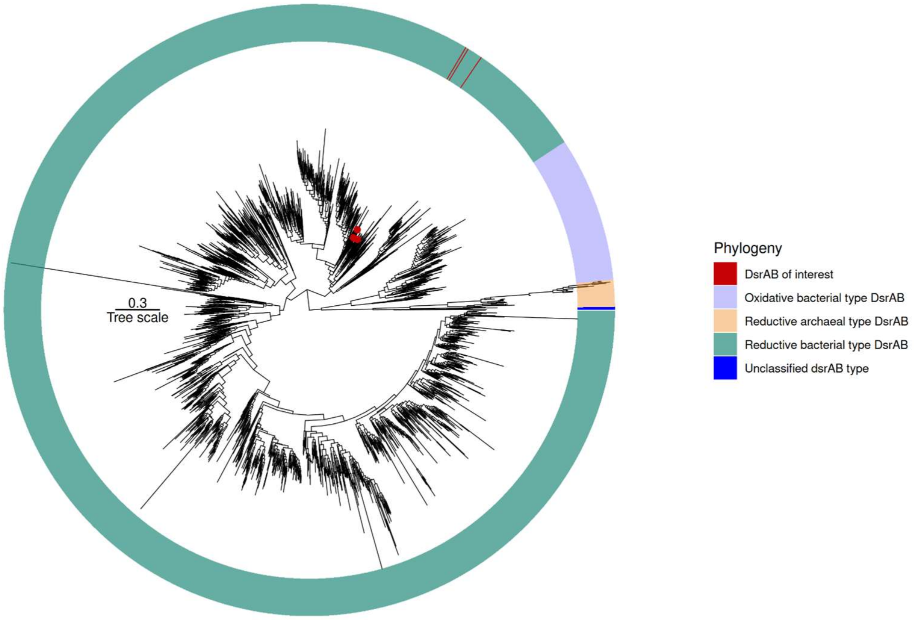


**Figure S8. Phylogenetic tree of DsrAB.** DsrAB from our Myxococcota MAGs are colored in red dots, and highlighted in red within the reductive bacterial type DsrAB.

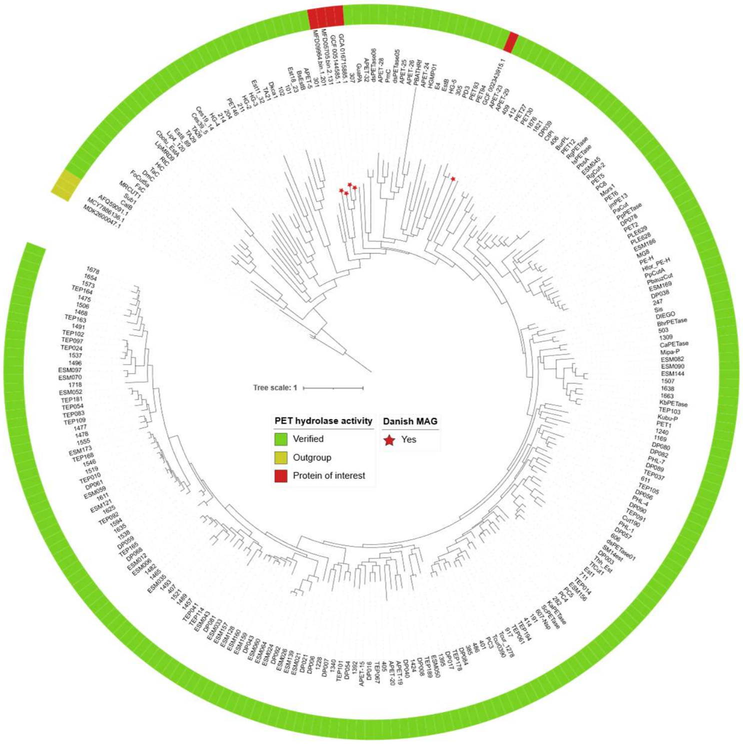


**Figure S9. Phylogenetic tree of functionally verified polyesterases having activity on PET.** PET hydrolases from our Myxococcota MAGs are labelled with a red star and highlighted in red in the outer ring. Functionally verified polyesterases having activity on PET downloaded from PAZy database [64] were in green in the outer ring with their enzyme names labelled. Putative PET hydrolase homologs from Danish MAGs clustered with experimentally characterized PET-active enzymes.
