## Supplementary material for "From soil to sea: unravelling the metabolic versatility and social dynamics of Myxococcota bacteria from different Danish environments": FigureS4

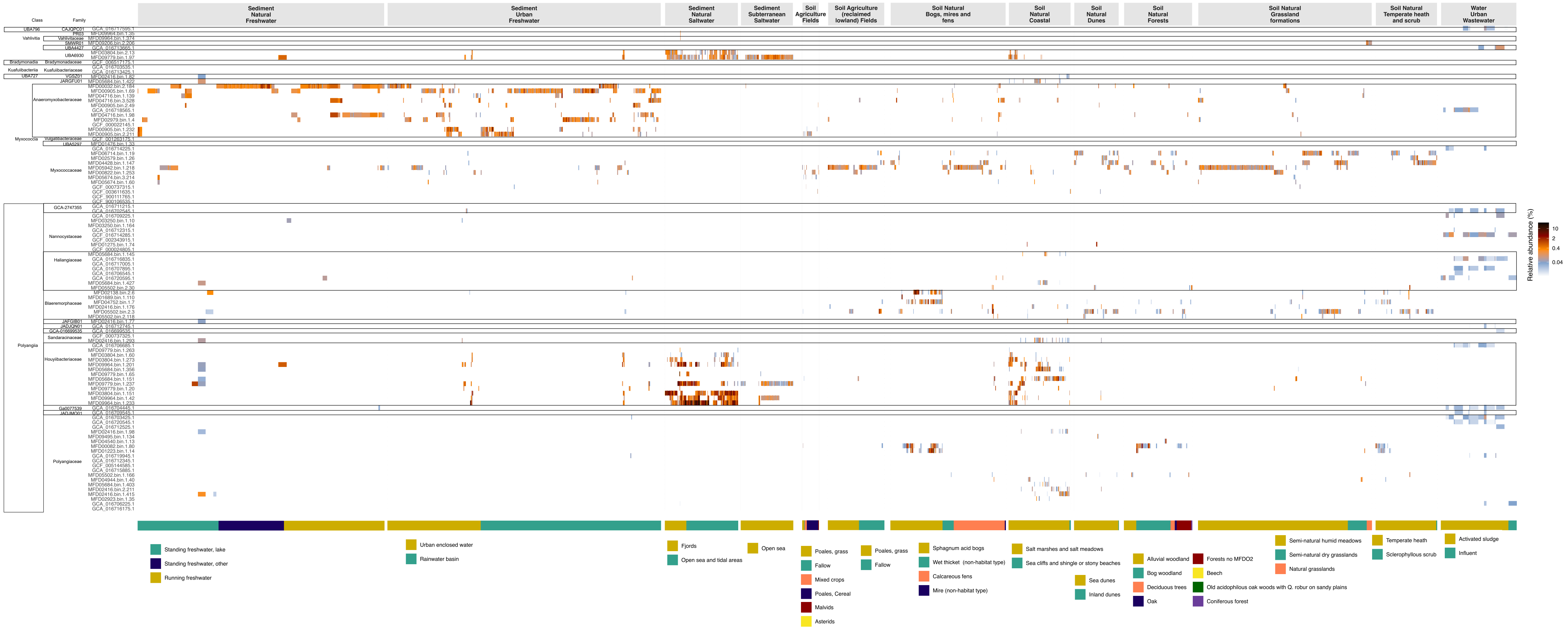

Class
Family
Sediment Natural Freshwater
Sediment Urban Freshwater
Sediment Natural Saltwater
Sediment Subterranean Saltwater
Soil Agriculture Fields
Soil Agriculture (reclaimed lowland) Fields
Soil Natural Bogs, mires and fens
Soil Natural Coastal
Soil Natural Dunes
Soil Natural Forests
Soil Natural Grassland formations
Soil Natural Temperate heath and scrub
Water Urban Wastewater
UBA796
Vahlivitia
Bradymonadia
Kuafuiibacteriia
UBA727
Myxococcia
Polyangiia
Vahlivitaceae
UBA6930
Bradymonadaceae
Kuafuiibacteriaceae
Anaeromyxobacteraceae
Vulgatibacteraceae
Myxococcaceae
GCA-2747355
Nannocystaceae
Haliangiaceae
Blaeremorphaceae
Sandaracinaceae
Houyiibacteriaceae
Polyangiaceae
Relative abundance (%)
10
2
0.4
0.04
Standing freshwater, lake
Standing freshwater, other
Running freshwater
Urban enclosed water
Rainwater basin
Fjords
Open sea and tidal areas
Open sea
Poales, grass
Fallow
Mixed crops
Poales, Cereal
Malvids
Asterids
Poales, grass
Fallow
Sphagnum acid bogs
Wet thicket (non-habitat type)
Calcareous fens
Mire (non-habitat type)
Salt marshes and salt meadows
Sea cliffs and shingle or stony beaches
Sea dunes
Inland dunes
Alluvial woodland
Bog woodland
Deciduous trees
Oak
Forests no MFDO2
Beech
Old acidophilous oak woods with Q. robur on sandy plains
Coniferous forest
Semi-natural humid meadows
Semi-natural dry grasslands
Natural grasslands
Temperate heath
Sclerophyllous scrub
Activated sludge
Influent
